# An automatic pipeline for the design of irreversible derivatives identifies a potent SARS-CoV-2 M^pro^ inhibitor

**DOI:** 10.1101/2020.09.21.299776

**Authors:** Daniel Zaidman, Paul Gehrtz, Mihajlo Filep, Daren Fearon, Jaime Prilusky, Shirly Duberstein, Galit Cohen, David Owen, Efrat Resnick, Claire Strain-Damerell, Petra Lukacik, Covid-Moonshot Consortium, Haim Barr, Martin A. Walsh, Frank von Delft, Nir London

## Abstract

Designing covalent inhibitors is a task of increasing importance in drug discovery. Efficiently designing irreversible inhibitors, though, remains challenging. Here, we present *covalentizer*, a computational pipeline for creating irreversible inhibitors based on complex structures of targets with known reversible binders. For each ligand, we create a custom-made focused library of covalent analogs. We use covalent docking, to dock these tailored covalent libraries and to find those that can bind covalently to a nearby cysteine while keeping some of the main interactions of the original molecule. We found ~11,000 cysteines in close proximity to a ligand across 8,386 protein-ligand complexes in the PDB. Of these, the protocol identified 1,553 structures with covalent predictions. In prospective evaluation against a panel of kinases, five out of nine predicted covalent inhibitors showed IC_50_ between 155 nM - 4.2 μM. Application of the protocol to an existing SARS-CoV-1 M^pro^ reversible inhibitor led to a new acrylamide inhibitor series with low micromolar IC_50_ against SARS-CoV-2 M^pro^. The docking prediction was validated by 11 co-crystal structures. This is a promising lead series for COVID-19 antivirals. Together these examples hint at the vast number of covalent inhibitors accessible through our protocol.

## Introduction

Covalent irreversible inhibitors have become increasingly popular over the last decade as chemical probes and drugs. Most often these inhibitors target a cysteine residue to form the covalent bond. Several rationally-designed irreversible inhibitors targeting cysteines were approved by the FDA in recent years, with notable examples such as Ibrutinib^1^, Afatinib^2^ and Osimertinib^3^. Irreversible binding offers a variety of advantages, including prolonged residence time^4^ and an ability to compete with high-affinity natural substrates^5–7^. Another important advantage of covalent inhibitors is their improved selectivity when targeting non-conserved cysteine residues^8,9^. Moreover, covalent binding can enable targeting of especially challenging targets such as the G12C oncogenic K-Ras mutation^10–12^.

Historically, most covalent inhibitors were designed by the addition of an electrophile to an already known reversible inhibitor that suitably binds next to a cysteine residue^13–18^. More recently, covalent inhibitors are also being discovered by empirical screening of covalent fragment libraries^19–25^ and by covalent virtual screening^10,26–32^. While covalent fragment and virtual screening can potentially discover new scaffolds, the binding affinity of primary hits may be relatively low, and often require laborious medicinal chemistry to reach suitable potency.

Covalent derivatization of an already known reversible binder, can endow the compound with added benefits of irreversible binding such as time dependent inhibition, longer duration of action, improved selectivity towards proteins that contain the target cysteine compared to homologs without a cysteine at that position, and possibly improved potency. Still, this approach is far from trivial. Three crucial questions have to be answered: 1. Which electrophilic moiety to use? 2. What is the optimal vector on the scaffold to attach through? 3. What linker, if any, would optimize the placement of the electrophile with respect to the binding mode of the scaffold and the position of the target cysteine residue? There are numerous possible answers for these questions. Furthermore, the *“covalentized”* (derivative containing the electrophile) version of the reversible inhibitor should be synthetically accessible. Therefore, tools that would enable to address this design problem algorithmically, would significantly simplify covalent inhibitor design and has the potential to discover many potent covalent binders for a large variety of targets.

Computational approaches to address this challenge are scarce. DUckCov^29^ a covalent virtual screening method, begins with non-covalent docking of a library of covalent compounds, while using pharmacophoric constraints for hydrogen bonds, as well as for the covalent warhead. This is followed by covalent docking of the ligands with the strongest non-covalent affinities. CovaDOTS^33^ uses a set of synthetic schemes and available building blocks, to create covalent analogs of existing non-covalent ligands, but was only assessed retrospectively. Cov_FB3D^34^ constructs *de novo* covalent ligands and was retrospectively assessed on recapitulation on known covalent inhibitors.

Here we present a computational pipeline to identify potential existing reversible binders for *covalentization* (creation of a covalent analog). Given a complex structure or model of a ligand in the vicinity of a cysteine residue, we elaborate the ligand or its substructures with various electrophiles. This *ad hoc* library of covalent analogs is covalently docked to the target protein and the original (non-covalent) structure is used as a filter to identify high-confidence covalent candidates. We applied this protocol - *covalentizer* - to the entire PDB to identify thousands of potential candidates amenable for irreversible inhibition, and made both the protocol and the database of pre-computed candidates publicly available to the community (https://covalentizer.weizmann.ac.il). We have prospectively synthesized and tested several predictions of various covalent kinase inhibitors proposed by the protocol and succeeded in five out of nine designs with IC_50_’s of 155 nM - 4.2 μM.

In early February 2020, the COVID-19 pandemic started to spread globally^35,36^. We turned to the pre-compiled database of *covalentizer* results, to look for possible candidate inhibitors for SARS-CoV-2 proteins. The search found a reversible small molecule inhibitor designed against the main protease of the SARS-CoV-1 virus (PDB: 3V3M^37^), which has 96% sequence identity to the main protease of SARS-CoV-2, with a promising covalent prediction. We synthesized the prediction and validated irreversible binding to the SARS-CoV-2 main protease (M^pro^). We further optimised the non-covalent affinity of the compound, resulting in improved analogs. Co-crystal structures confirmed the computational model. This example highlights the strength of our method - the design was already available, and enabled very rapid development. The database suggests that hundreds more such examples await testing.

## Results

### The covalentizer pipeline

For a given complex structure with a reversible ligand in the vicinity of a target cysteine residue, the pipeline (Fig. 1) comprises four consecutive steps: fragmentation, electrophile diversification, covalent docking and RMSD filtering.

**Figure 1.**
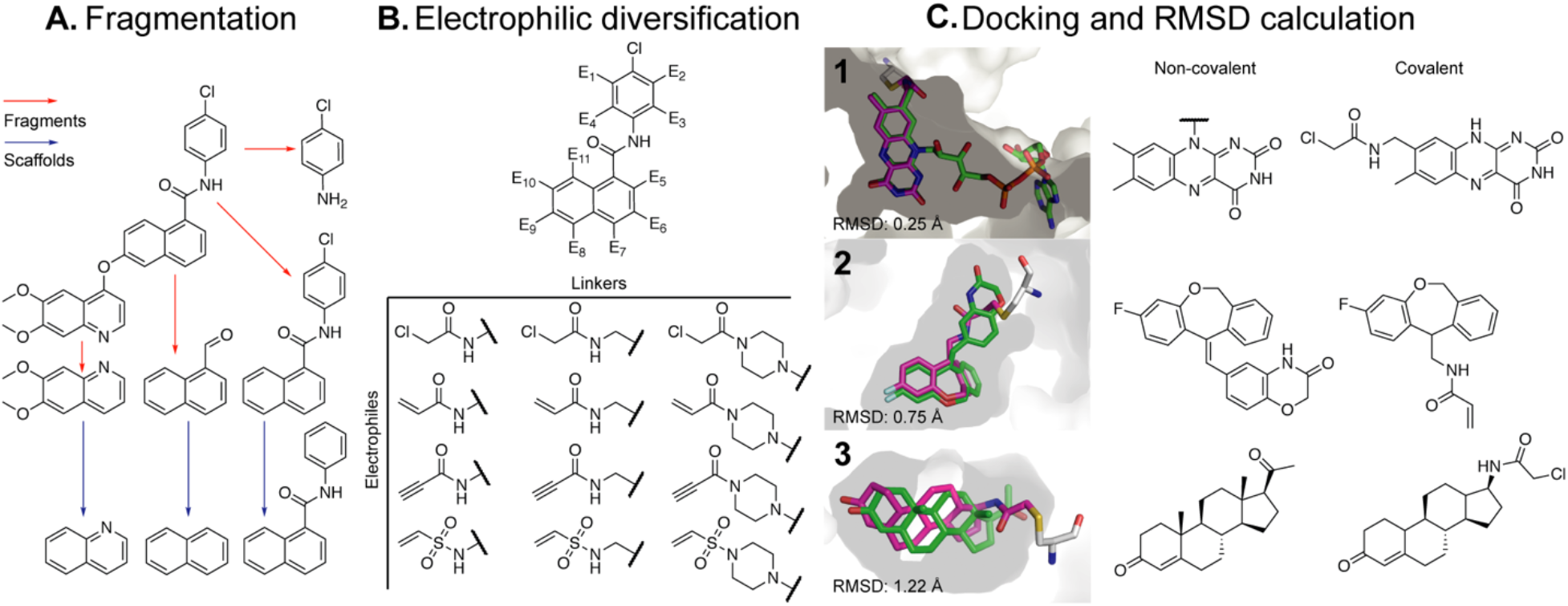
An overview of the *covalentizer* computational protocol. The protocol comprises four consecutive steps. **A.** Fragmentation: the molecule is broken and divided into fragments (red arrows) using synthetically accessible bonds^38^. Murcko scaffolds^39^ of the fragments (blue arrows) are also added to the list of fragments. **B.** Electrophilic diversification: for each substructure, a library of potential electrophilic analogs is generated, a few hundred compounds in size. We used four kinds of nitrogen-based electrophiles ranging in reactivity: vinyl sulfones, chloroacetamides, acrylamides and propynamides. We also considered various linkers between the fragment and the electrophile. **C.** Docking: The target structure is then docked against its appropriate analog library using all available cysteine rotamers. Finally, RMSD calculation: For each docked compound, an RMSD is calculated between the MCS (maximal common substructure) of the reversible compound and the new covalent analog. We show examples of predictions with increasing RMSDs, for binders of 1. Nitrate reductase from Ulva prolifera (PDB: 5YLY) 2. Human mineralocorticoid receptor (PDB: 5HCV) and 3. Human progesterone receptor (PDB: 1A28).

#### A. Fragmentation

In this step, the ligand is broken-down and divided into two parts via synthetically accessible bonds^38^. Doing this recursively, results in a list of substructures (Fig. 1A). For each substructure, we augment the list with its corresponding Murcko scaffold^39^ (the naked ring system, without any decoration) to allow more exit vectors from which the electrophile can be added next. The motivation for this fragmentation step is three-fold. First, as mentioned, fragmenting the molecule exposes new vectors on which to install the electrophile (see Fig. 1C example 2). Second, the additional constraint of forming the covalent bond might cause a slight shift to the molecule’s binding mode from the original crystal structure. Such a shift may propagate and cause a steric clash between the protein and a ligand moiety distal to the electrophile. Since adding the covalent bond is expected to increase the overall potency, we ‘sacrifice’ parts of the molecule to enable the addition of an electrophile. The final ranking of candidate covalent analogs relies on covalent docking which is sensitive to sub-Å shifts. Hence, occasionally, a truncated version of the ligand will dock well, while the full ligand will not. Thus, including the sub-structures and their scaffolds maximizes the number of candidates. Since covalent docking accuracy was shown to deteriorate with ligand size and number of rotatable bonds^26^, we filter the final list of substructures, to those with 8-25 heavy atoms and up to five rotatable bonds.

#### B. Electrophile diversification

For each substructure or scaffold, we generate a library of potential electrophilic analogs, typically resulting in a few hundred analogs (Fig. 1B). We consider four kinds of electrophiles ranging in reactivity: vinyl sulfonamides, chloroacetamides, acrylamides and propynamides, that can all be installed in one step onto a free amine. We add these electrophiles to the substructures using simple connection rules which, however, do not guarantee synthetic accessibility (see methods section for more details). We also consider various linkers between the fragment and the electrophile. In our application below, we considered either a methylene linker or various di-amine linkers (Supp. Fig. 1).

#### C. Covalent docking

The structure of the complex is prepared for docking, using all available cysteine rotamers. We use DOCKovalent^26^ to dock the electrophile library we described above against the protein (after removing the crystallographic ligand).

#### D. RMSD filtering

Compounds that are able to form a covalent bond with the target cysteine while still maintaining the same binding mode of the original reversible ligand are likely candidates for covalent analogs. To assess this, we evaluate the RMSD (root mean square deviation) between each docking prediction and the crystallographic ligand. Due to the fact that RMSD is calculated between two sets of matching atoms, and the reversible ligand is different from the irreversible one, it was calculated based on the maximal common substructure (MCS) between the two molecules. Figure 1C exemplifies predictions with varying RMSDs. For a PDB wide application of the pipeline, we focused on covalent analogs with a docking position of < 1.5 Å RMSD from the crystallographic ligand.

### Covalent kinase inhibitors benchmark

To benchmark the pipeline, we wanted to test whether it is able to find known covalent inhibitors, given only their reversible part as input. To achieve this, we used the kinase subset of a recently published covalent docking benchmark^30^. This set included 35 kinase covalent inhibitor complex structures with either acrylamides, chloroacetamides or vinyl sulfonamides (after excluding seven inhibitors with uncommon electrophiles). To form the input for *covalentizer*, we removed the electrophiles while leaving only a free amine. For substituted acrylamides we removed β-substitutions as well.

Out of the 35 structures, the pipeline identified the crystallographic covalent inhisbitor in 14 (40%) of the cases, with a threshold of 1.5 Å MCS-RMSD (Fig. 2; Supp. Table 1).

**Figure 2.**
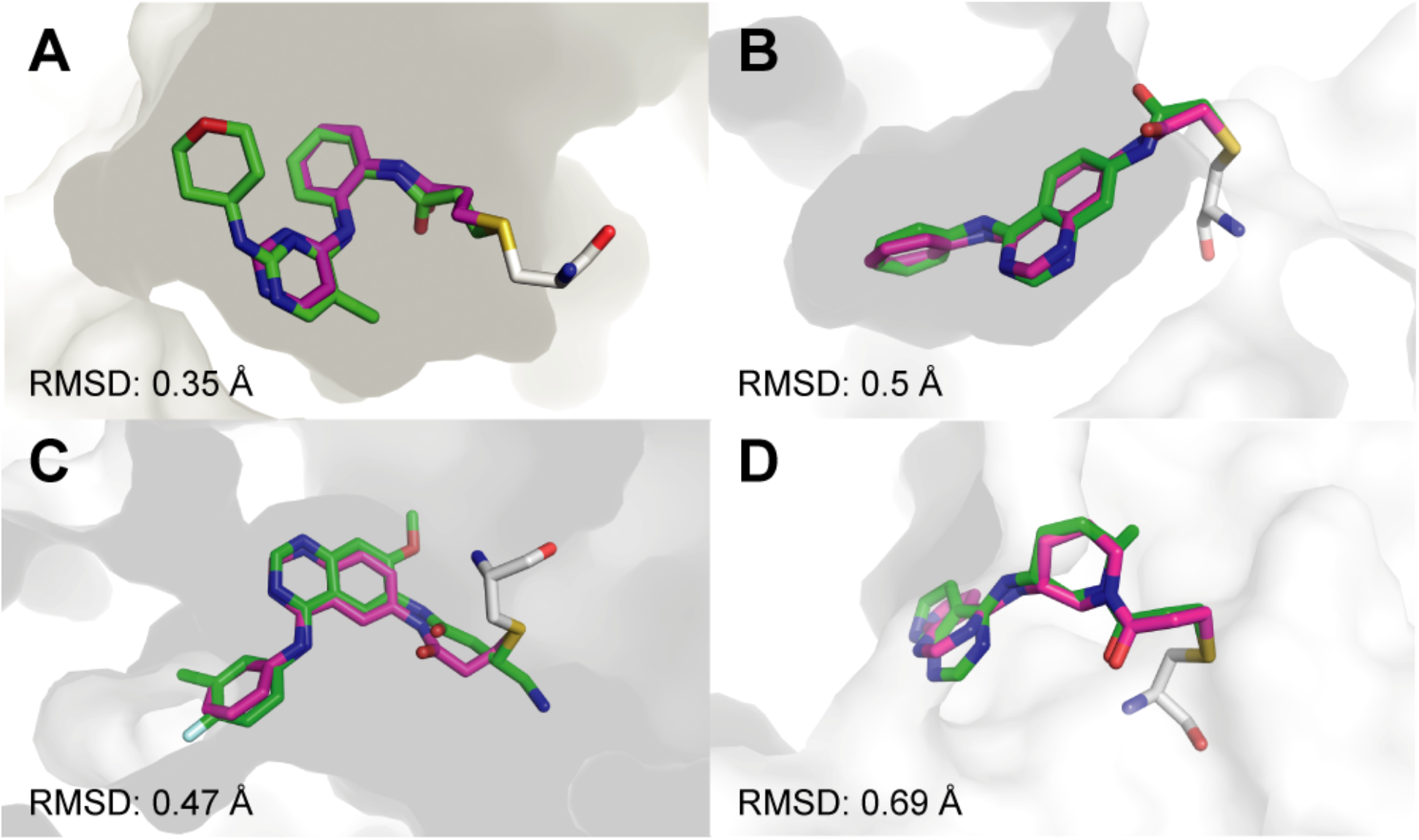
*Covalentizer* successfully recapitulates known covalent kinase inhibitors. Examples of covalent kinase inhibitors (green) for which *covalentizer* was able to find a substructure match (magenta) under the 1.5 Å threshold. **A.** ERK2, PDB: 4ZZO. **B.** EphB3, PDB: 5L6P. **C.** EGFR (T790M), PDB: 4I24. **D.** JAK3, PDB: 5TOZ. The electrophiles span acrylamides (A,D), a substituted acrylamide (C) and chloroacetamide (B).

### Covalentizing the PDB

Encouraged by the results in recapitulating known covalent kinase inhibitors we aimed to apply our protocol to the entire PDB. We started from the set of all the protein-small molecule X-ray structures (< 3.0 Å resolution) that contained a small molecule with a molecular weight greater than 300 Da, and no DNA/RNA chains. As of the date of the search (July 4th, 2019) this resulted in 44,990 structures. We filtered these to structures in which a ligand has one of its atoms within 6 Å of the sulphur atom of a free cysteine residue. Disulfides or covalently modified cysteines were excluded. After applying this filter, we ended up with 8,386 such structures, and ~11,000 cysteines.

These structures, which constitute the target space for our protocol, contain significant redundancy. Clustering them with a threshold of 90% sequence identity, results in 2,227 representatives. 38% of the structures are of human proteins and the rest span many other organisms including rodents, bacteria and yeast. They also span seven different enzyme classes, with the most prevalent being transferases (41.4%). 928 structures (11.1% of the entire dataset) are kinases. These ~8,400 proteins contain 3,673 different ligands, each binding next to a cysteine. The ligand that is most abundant in this database is Flavin-Adenine Dinucleotide occurring in 504 structures, whereas 3,058 ligands (83% of the compounds) occur only in a single structure. The most common ligands were nucleotide or nucleotide-like molecules.

After running the aforementioned algorithm against the ~8,400 structures that passed our filtering (Fig 3A), 1,553 structures produced at least one candidate below the 1.5 Å RMSD cutoff. These structures represent roughly 380 proteins (representative set at 95% sequence identity). 1,051 structures are of human proteins, 338 are structures of kinases. 80 of the structures had produced a covalent analog prediction that was docked <0.5 Å from the original ligand, representing very high-confidence candidates (Fig. 3B). The distribution of selected electrophiles is almost uniform (Fig. 3C). All of the predictions are made available through a public website (https://covalentizer.weizmann.ac.il) which is automatically updated weekly with new PDB entries.

**Figure 3.**
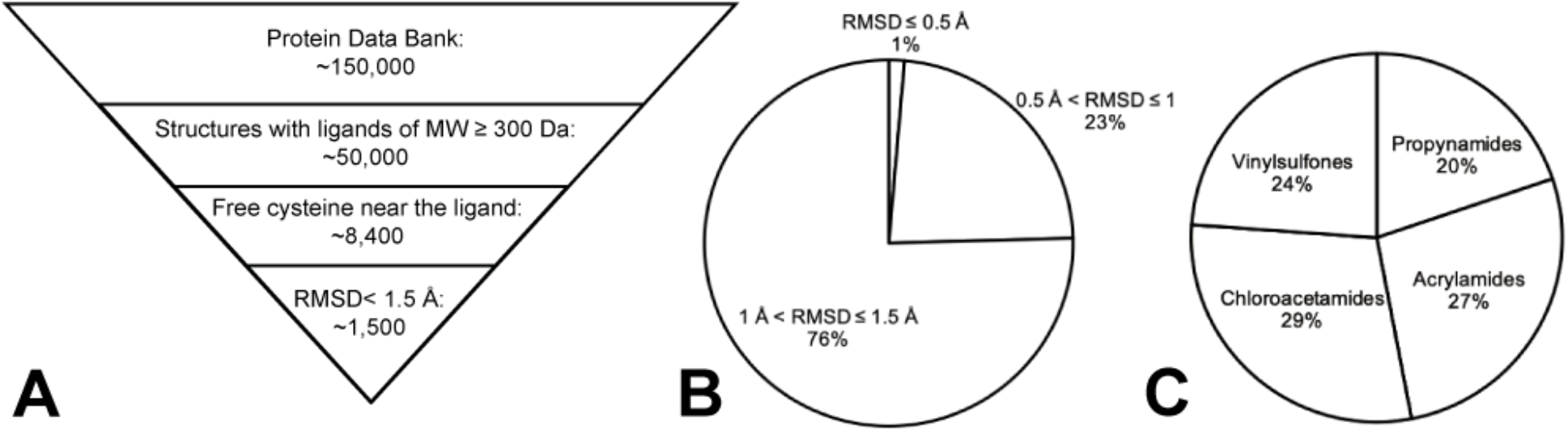
PDB wide application of *covalentizer* identifies candidate irreversible inhibitors for more than 1,500 structures. **A.** We filtered the protein data bank (PDB) for structures that had only protein chains (no DNA/RNA), and contained a small molecule of at least 300 Da. This threshold was set to ensure some minimal initial fit/binding affinity to the target, as well as to filter out non-ligand small molecules like crystallization reagents. We used a pymol based script to filter only the structures in which at least one ligand atom is < 6 Å away from the sulfur atom of a cysteine residue. This cysteine also has to be free (no disulfide or other covalent modifications). After running the *covalentizer* protocol and filtering only for results with < 1.5 Å RMSD of the maximal common substructure (MCS) between the reversible ligand and the covalent analog generated by *covalentizer*, there were 1,553 structures for which at least one such prediction was obtained. **B.** The top 1% of results have an RMSD under 0.5 Å. 23% are between 0.5 Å and 1 Å, and 76% are between 1 Å and 1.5 Å. **C.** The distribution of the four electrophiles used is balanced, with 29% chloroacetamides, 27% acrylamides, 24% vinylsulfones, and 20% propynamides.

### Exploring additional linkers

As mentioned above, the entire database was processed using direct attachments of the electrophiles to atoms of the sub-structures, as well as with a methylene linker. The use of longer and more diverse linkers for the addition of an electrophile would allow the targeting of cysteines further from the ligand thus increasing the available target space, as well as diversifying the introduced chemistry. To investigate this further, we searched the covalent inhibitor discovery literature^40–43^ for the most common di-amine linkers used in the last decade which led to the selection of 7 aromatic linkers and 17 aliphatic linkers (Supp. Fig. 1). Since including all of these linkers increases the computational demands of the pipeline, we restricted its application to the subset of liganded kinase structures in the PDB. Since these linkers can enable ligands to reach further cysteines at extended distances, the search criteria was extended to a distance of up to 10 Å from the ligand (instead of 6 Å, previously).

The final subset includes 1,880 PDB structures that contain a Cys residue of up to 10 Å away from one of 1,398 various ligands. The size of the custom-made libraries of electrophilic analogs for a particular reversible ligand, containing these linkers, now extends to a few thousand compounds. Overall, we generated *in silico* over 3 million electrophilic compounds with di-amine linkers for the kinase subset. The results show candidates of < 1.5 Å MCS-RMSD between the original reversible ligand and the electrophilic candidate for 411 protein structures. 186 of these structures (45%) were not found in the previous run, showing the potential of using these more sophisticated linkers to reach farther cysteines and to *covalentize* more ligands.

### Novel covalent inhibitors for various kinases

Kinase inhibitors comprise 22% of the *covalentizer* results. We selected a subset of these for prospective validation. We chose the candidates based on three features: 1. Low RMSD relative to the parent reversible ligand. 2. The addition of the electrophile is not predicted to interfere with the kinase hinge binding region. 3. Ease of synthesis. This required manual inspection of pre-selected low RMSD results. Overall, we made and tested nine compounds (Fig. 4) targeting five different kinases. In some cases, addition of the electrophile required removal of large parts of the parent reversible ligand (Supp. Fig. 2). The compounds were each tested in a kinase activity assay against the target kinase in the structure from which the *covalentizer* result was derived. The assay was performed at ATP concentration equal to the Km of the kinase in question, with a 2 h pre-incubation of the inhibitor at 25 °C.

**Figure 4.**
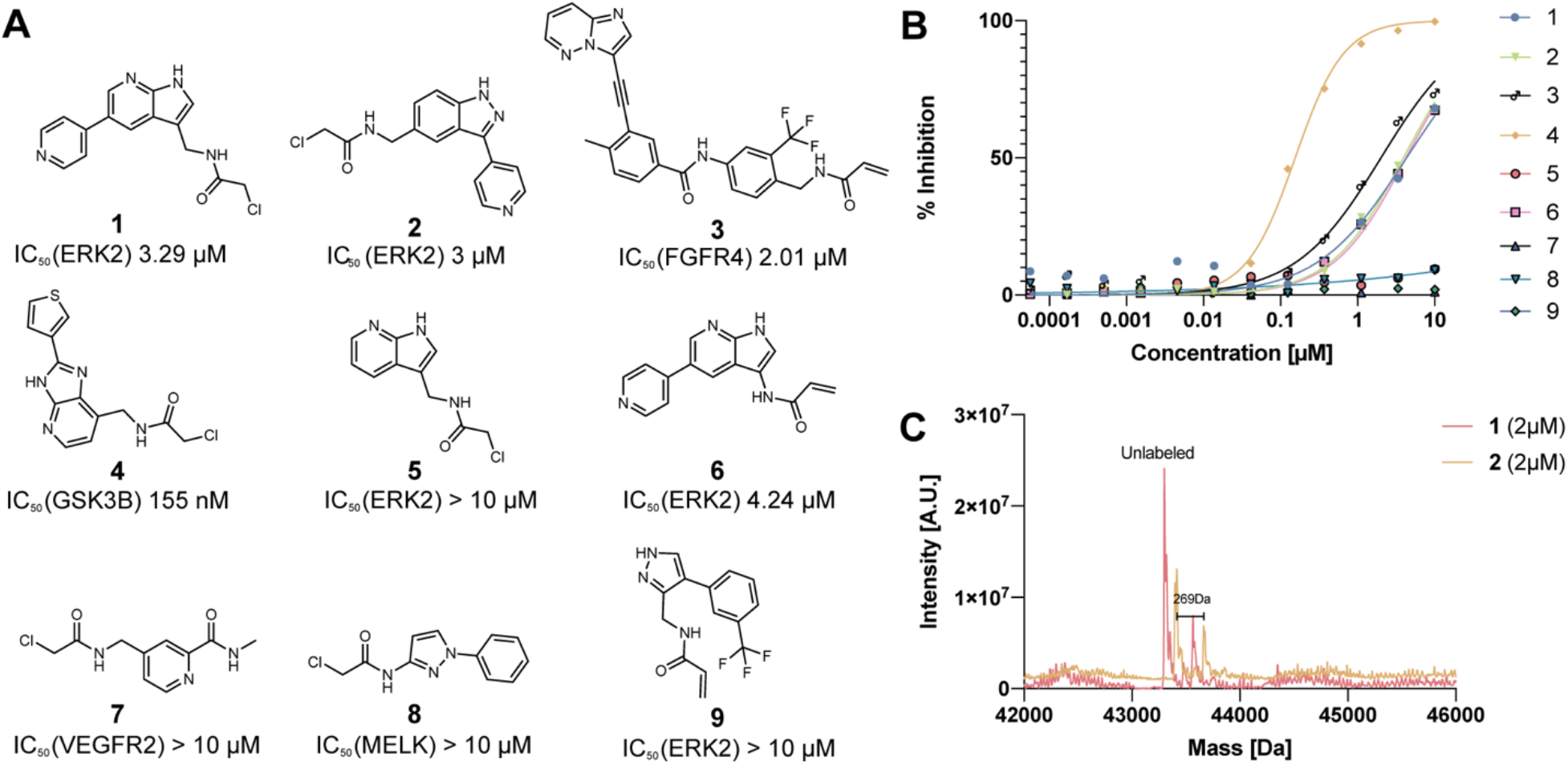
Prospective prediction identifies novel irreversible kinase inhibitors. **A.** Chemical structures and in vitro kinase activity assay IC_50s_ for nine prospective *covalentizer* predictions. See Supp. Fig. 2 for the parent compounds, pose predictions and RMSD values. **B.** Dose response curves for each of the nine compounds (see additional repetitions for **1** and **2** in Supp. Fig. 3). Each compound was tested against its corresponding target kinase. **C.** Deconvoluted mass spectra obtained by intact protein LC/MS of recombinant ERK2 (2 μM) incubated with equimolar **1** or **2** for 1 h at room temperature, identifies significant irreversible binding by both compounds.

Four of the nine compounds did not show inhibition under the assay conditions (IC_50_ > 10 μM). Three compounds targeting ERK2 showed IC_50_ values of 3 - 4.24 μM. For two of these inhibitors, **1** and **2**, we assessed irreversible binding to ERK2 by intact protein mass spectrometry (2 μM ERK2, 2 μM compound, 1 h incubation at 25 °C). The expected protein-compound adducts were detected (25% and 33% labeling respectively; peak-to-peak Δm 265-270 Da for both compounds; Fig. 4B) with no additional adducts derived from multiple reactions, highlighting the moderate reactivity of the designed α-chloroacetamides. The remaining *covalentizer* hits included a 2.01 μM inhibitor (**3**) of FGFR4 derived from the non-selective kinase inhibitor ponatinib, and a 155 nM inhibitor (**4**) of GSK3β.

### A covalent SARS-CoV-2 main protease inhibitor

Upon the release of the first structure of the new SARS-CoV-2 M^pro^ protease (PDB: 6LU7^44^) we noticed that the active site is nearly identical to that of SARS-CoV-1. The entire protein is highly conserved with 96% sequence identity. This prompted us to search the database for covalent versions of SARS-CoV-1 M^pro^ ligands. One such prediction was available based on a reversible inhibitor ML188 (IC_50_= 4.8 ± 0.8 μM, racemate; 1.5 ± 0.3 μM, (*R*)-enantiomer) of the SARS-CoV-1 main protease (PDB: 3V3M^37^; Fig. 5A). We re-synthesized and tested racemic ML188 against SARS-CoV-2 M^pro^ which showed an IC_50_ of 3.14 μM (Supp. Fig. 4A), similar to what has been reported for SARS-CoV-1. ML188 was synthesized using the Ugi four-component reaction (4-CR), and the covalent prediction was easily accessible by replacing one reactant (2-furoic acid to acrylic acid) to give **10**, synthesized and isolated as the racemate (Fig. 5D). We initially assessed irreversible binding of **10** towards recombinant SARS-CoV-2 M^pro^ using intact protein mass spectrometry (2 μM protein, 1.5 h incubation with electrophile at 25 °C; Fig. 5F). The expected adduct was detected with 19% labeling at 2 μM compound, and up to 88% labeling at 200 μM compound (Fig. 5F).

**Figure 5.**
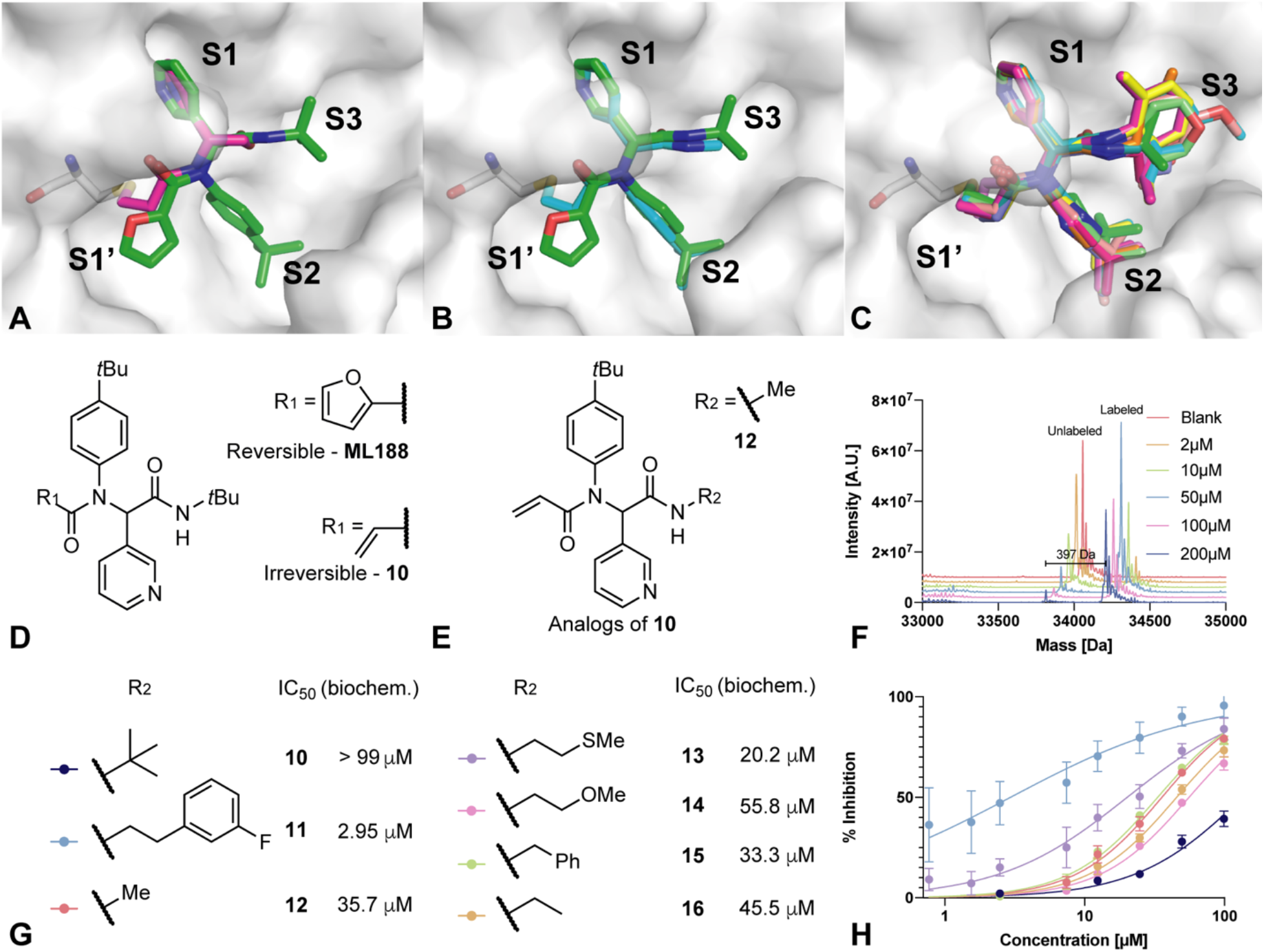
Computational prediction and experimental validation of an irreversible SARS-CoV-2 M^pro^ inhibitor. **A.** The *covalentizer* prediction of **10** (magenta) overlaid on the non-covalent compound it is based on (ML188^45^; green; PDB:3V3M). The protocol suggested to substitute the furanyl moiety of ML188 with an acrylamide to bind the catalytic cysteine. The RMSD between the covalent fragment and the original reversible inhibitor is 0.65 Å. **B.** The crystal structure of one of the covalent analogs of **10** (PDB: 5RH5; cyan) overlaid on ML188 (green). **C.** Overlay of all the 11 crystal structures of compound **10** analogs, all exhibiting the same predicted binding mode. PDBs: 5RGT, 5RH5, 5RH6, 5RH7, 5RH9, 5RL0, 5RL1, 5RL2, 5RL3, 5RL4, 5RL5. For individual structures see Supp. Fig. 5. **D.** The chemical structures of ML188 and **10**. **E.** Chemical structure of Ugi compounds exploring the S3 pocket, with the R group that is shown in the crystal structure in (B). **F.** Deconvoluted mass spectra obtained by intact protein LC/MS of recombinant SARS-CoV-2 M^pro^ 2μM incubated with 2 μM - 200 μM **10** for 1.5 h at room temperature. **G.** Further analogs of **10** with their associated biochemical potencies. **H.** The dose response curves for the seven compounds shown in G.

Despite the irreversible binding, this initial compound did not show strong inhibition in a fluorescence-based enzymatic assay (IC_50_ > 99 μM, 13% inhibition at 20 μM; 15 min pre-incubation; Fig. 5H). However, it was a promising starting point for additional optimization. Due to the modular nature of the Ugi 4-CR procedure, it was possible to synthesize and test large libraries of analogs by systematically varying each reactant to target different pockets. We designed those libraries based on computational modelling of *in silico* generated Ugi products, as well as an exhaustive screen of commercially available isocyanides (see Supp. Dataset 1). A few of the early combinatorial synthetic results, which had low biochemical potency (comparable to our starting Ugi compound), allowed for crystallographic analysis in the presence of M^pro^. In these cases, the expected binding mode was recapitulated experimentally and showed low deviation from the non-covalent starting point (Fig. 5B, 5C, Supp. Fig. 5), thus proving the *covalentizer* prediction to be correct. In all crystal structures, the electrophile formed the expected covalent bond with the catalytic cysteine residue.

To optimize **10**, we have made and tested close to 140 analogs (Supp. Dataset 1; Fig. 5. Supp. Fig. 7,8), exploring all three components of the Ugi reaction while keeping the acrylamide fixed. We explored a variety of replacements for the initial *p-tert*-butylphenyl motif protruding into the S2 pocket (Fig. 5), most of them did not result in improved potency (Supp. Fig. 7C). Similarly, independent optimization of binding to the S1 pocket only led to the identification of one beneficial change (**23**, IC_50_ 65.58 μM), with a *meta* chloro-substitution of the pyridine (Supp. Fig. 7A). Other substituents (-Br, -OMe, -OEt, -CF_2_CH_3_) led to inactive compounds.

Beyond further optimization of the S1 and S2 pocket binding it was clear that extension of the ligand towards the S3 and S4 pockets should prove fruitful. For example, a reversible-covalent α-ketoamide inhibitor^46^ (biochemical IC_50_ 0.67 μM ± 0.18 μM) probes the S3/4 region with an additional hydrogen bond to the backbone of Glu166. In a large scale fragment screen, numerous fragments were able to bind in these pockets^47^. In this case, we exhaustively synthesized analogs of **10**, using 34 available isocyanides. Starting from **10**, simple alkyl chain extension resulted in compounds with improved potency (Fig. 5D). In particular compound **11**, harboring a phenethylamide motif, was particularly potent with an IC_50_ of 2.95 μM (Fig. 5G) and K_inact_/K_i_ of 18.4 M^−1^s^−1^ (Supp. Fig. 6).

It appears that relief of steric strain around the amide nitrogen also plays a part, since change to a methyl amide (in **12**, relative to **10**) also resulted in increased potency. Opposed to our initial assumption of independent optimization of S1-3 pocket binding, the combination of beneficial structural motifs in a third generation of Ugi products led to inhibitors with diminished potency compared to **11** (see Supp. Fig. 7B; Supp. Dataset 1). One explanation for this behaviour is the high plasticity of M^pro^, leading to induced fit effects.

Removal of the furanyl in ML188 and replacement with an electrophile in **10** initially led to a loss in potency, which in this case was overcome by optimization of the non-covalent affinity in the S3 region to give compound **11**. Re-installing the furanyl ring in combination with the S3-optimized phenethylamide motif led to compound **17** with a similar IC_50_ (2.72. μM; Supp. Fig. 4B), suggesting that the marked improvement of this side-chain is particular to the covalently bound conformation.

In conclusion, we successfully executed a mode of action change towards irreversible targeting of the catalytic cysteine residue in M^pro^ which may have improved activity in cells as well as long-term strategic benefits to safeguard against viral evolution.

## Discussion

Designing new covalent inhibitors is challenging. Here we leveraged the subset of protein targets for which a structure of a known binder is available, to computationally enumerate and evaluate exhaustive sets of covalent derivatives. Automating the protocol allowed us to apply it to the entire PDB and assess the applicability of this approach. Prospective testing against six real-world targets demonstrated that irreversible ligands can be reached with little synthesis, and structures validated the binding-pose prediction.

A main advantage of our work is the wide exploration of X-ray structures which produced an extensive list of candidates waiting to be explored. This allowed us to quickly find a promising lead series against SARS-CoV-2 M^pro^. This prediction, which was based on a historic non-covalent SARS-CoV-1 M^pro^ inhibitor^45^ was pre-calculated, and ready for synthesis at a moment’s notice. We have made thousands of such predictions available through a public website (https://covalentizer.weizmann.ac.il) which updates weekly with the release of new structures to the PDB. It also allows *covalentizing* of user uploaded structures. We believe this would enable wide application and experimental testing of new covalent inhibitors.

Despite the success of our protocol, several caveats remain. First is the fact that currently the protocol does not take into account the synthetic feasibility of the proposed designs. When selecting candidates for prospective evaluation, we found that some of the molecules required complicated synthesis. Incorporating into our pipeline a strategy such as DOTS^33,48^, other retrosynthesis algorithms^49–51^ or even the use of synthetic feasibility scores^52–54^, can significantly improve the quality of proposed candidates in the future.

Another point for improvement is the relatively weak potency of our prospective designs in comparison to their parent compounds. One likely explanation for these lower affinities is the removal of non-covalent affinity elements which are not sufficiently compensated by the gains from covalent bond formation. For example, in compound **2** (derived from PDB: 4QTA) more than 350 Da of the original compound^55^ is removed (Supp. Fig. 2), resulting in three orders of magnitude loss in potency. However the remaining covalent fragment still shows significant inhibition of ERK2. Another example is compound **1**, its parent compound (PDB: 4QP9) has an IC_50_ of 71 nM, however the propyl-pyrazole group we have omitted in order to accommodate the electrophile (Supp. Fig. 2) improved the parent reversible binder by more than 150-fold. Lastly the loss of a hydrogen bond between the M^pro^ backbone NH of Gly143 and the furanyl oxygen of ML188 (PDB: 3V3M), decreased potency, under our assay conditions of 15 min. pre-incubation, by more than 30-fold. These results suggest that a careful examination of the binding energy contribution is required for the parts that are omitted in order to accommodate the new electrophile.

However, as we saw for both ERK2 (**1** *vs*. **5**) and M^pro^ (**10** *vs*. **11**), improving reversible recognition is able to improve potency, and even to surpass the parent compound in the case of **11**. For this series we believe that further structure-based optimization of binding to the S3 pocket, including H-bonding to Glu166 and chiral separation of the active enantiomer can pave the way for sub-micromolar Ugi-type covalent M^pro^ inhibitors. Thus, in many cases where irreversible binding is needed our protocol can provide a promising starting point for optimization.

Another possible explanation for the relatively low affinity of the irreversible binders are slight inaccuracies in the covalent warhead positioning which results in sub-optimal covalent bond formation. Perhaps due to the fact that the docking program does not take into account the actual formation of the covalent bond, and ignores for instance the transition state energy of the rate-determining step of the organosulfide bond formation, but rather evaluates the binding energy of the adduct. Better understanding of the steric and electronic constraints of the covalent bond formation, and hence a better docking software should improve the results.

The docking software also ignores the intrinsic reactivity of the proposed designs. It is interesting to note in this regard the similar activity of a methylene-chloroacetamide (**1**), compared to its acrylamide analog (**6**). Such electrophile replacements can be very useful in rational design of irreversible inhibitors, especially if they prove to work across various scaffolds. Geometrically, the additional methylene before the chloroacetamide makes the distance from the ring to the thiol similar to that of the acrylamide (Supp. Fig. 2). In terms of reactivity, however, the acrylamide, conjugated to the azaindole is activated^56^ and thus is likely closer in reactivity to the chloroacetamide. Indeed, a methylene linker would be the minimal linker element required to insulate against π-conjugation, allowing easier prediction of intrinsic reactivity. No-linker designs connected to extended π-systems such as heteroarenes often exhibit a range of intrinsic reactivities^20,56^ which remain challenging to predict computationally^57–59^ and thus require careful evaluation.

Many additional designs remain to be discovered beyond the more than 1,500 we made available through the *covalentizer* server. New electrophiles and linkers which will enable new geometric trajectories between the cysteine and the molecule, can considerably expand the design space. We tested this idea computationally using a library of linkers curated from the literature (Supp. Fig 1), on a subset of kinases from our database, showing an increase in the number of structures that can be *covalentized*. New covalent ‘warheads’, including reversible covalent warheads, such as cyanoacrylamides^60^, and clorofluoroacetamides^61^ become available, both for cysteine residues^62^, but also for other amino acids^63–65^. These can be incorporated with little effort into the *covalentizer* pipeline. Since cysteine is one of the least abundant natural amino acids, additional covalent chemistries will significantly expand the number of ligands that can be potentially addressed.

In summary, we show that using covalent docking we were able to make irreversible analogs of ligands for which a complex structure is available. We made our discoveries public in the form of a database of the results we obtained by running our protocol on the entire PDB which is automatically updated weekly with newly released entries, as well as a web-tool for applying the protocol on new targets given by users. Using the protocol, we discovered new covalent kinase inhibitors and optimised a potent covalent COVID-19 protease inhibitor, with a low-cost, modular and fast synthesis. We hope our results will encourage researchers to apply covalent inhibitors for a wide range of targets.

## Methods

### Programs and libraries

RDKit was used for 2D molecular handling, conformation generation and RMSD calculation. RDKit: Open-source cheminformatics; RDKit.org. Marvin was used in the process of preparing the molecules for docking, Marvin 17.21.0, ChemAxon (https://www.chemaxon.com). DOCKovalent ^26^ was used for virtual covalent docking.

### Curating target structures from the PDB

Using pymol scripts (The PyMOL Molecular Graphics System, Version 2.0.4 Schrödinger, LLC), we filtered only the structures that have a ligand in which one of its atoms is within 6 Å cysteine residue. We further filtered the list to include only cysteines with a free thiol group (defined as a sulfur atom that is only connected to the residue’s Cβ). By doing this, we discarded any disulfides, as well as cysteines that are already covalently attached to a ligand. We further removed any ligands which had more than one copy per chain in the structure, and ligands on which processing of the ligand’s SMILES failed.

### Enumerating substructures for covalentization

Fragmentation and scaffold extraction was done using RDKit’s implementation of the Recap algorithm ^38^ and the MurckoScaffold ^39^ functionality respectively.

### Covalentizing a substructure

For each substructure or scaffold, we generated a library of potential electrophilic analogs using SMARTS based reactions. The reaction rules were: 1. Adding an electrophile (including the nitrogen) to any non-substituted aromatic carbon, as well as all aliphatic carbons with one or two bonded atoms, excluding carbons which are already connected to nitrogen. 2. Adding an electrophile to a free amine, either primary or secondary. In this case the nitrogen is completed with the rest of the electrophile. The first rule will usually require more complicated synthesis, whereas the second rule, will allow to use the same ligand as a starting material for a nucleophilic substitution of the acyl form of the electrophile with the free amine.

### Docking and RMSD calculation

RDKit and Marvin were used to create 250 conformations for each electrophilic analog. Covalent docking was done using DOCKovalent – a virtual screening program. We docked the appropriate analog library for each target, while saving 10 structures for each analog to increase the number of final candidates. When docking the larger linker based libraries we only used the top scoring structure for each analog, due to the large number of structures for analysis. Alternative rotamers for the cysteine residue were generated with pymol based scripts. We used RDKit to filter only for results with a MCS that has an RMSD of less than 1.5 Å to the original ligand.

### Computational optimisation of the M^pro^ inhibitor

We used the RDKit reaction functionalities, as well as OpenBabel (http://openbabel.org/wiki/Main_Page) to prepare virtual libraries of analogs of compound **10**. The Ugi reaction has three reactants: amine, isocyanide, aldehyde and carboxylic acid. The carboxylic acid is set constant to acrylic acid, since we didn’t want to change the electrophilic component. In the virtual libraries, we left it as the reversible furan moiety for convenience in modeling. We thus created three such libraries, each one by replacing one of the three other Ugi reactants with commercially available building blocks. Using RDKit, we generated up to 100 constrained conformations of each molecule, by fixing the conformation of three components as in the crystal structure, and changing only the conformation of the variable part. We then used the Rosetta modeling suite in order to choose the best conformation for each compound, when bound to the protease. For each molecule, we then defined this set of constrained conformations as an extra residue for Rosetta, and used Rosetta Packer^66^ to choose the best conformation, while allowing side-chain flexibility. Eventually, we chose analogs only for the amine and the isocyanide components, as the aldehyde component was highly optimised already. We chose 9 isocyanide replacements and 14 amine replacements (one of them was not based on docking). Most combinations of these components were made by Enamine and tested as part of the Covid-Moonshot effort^47,67^.

### Intact protein LC/MS

M^pro^ was incubated for 90 minutes in 50 mM Tris pH 8 300 mM NaCl in room temperature. ERK2 was incubated for 60 minutes in 10 mM Hepes pH 7.5 500 mM NaCl and 5% glycerol in room temperature. The LC/MS runs were performed on a Waters ACUITY UPLC class H instrument, in positive ion mode using electrospray ionization. UPLC separation used a C4 column (300 Å, 1.7 μm, 21 mm × 100 mm). The column was held at 40 °C and the autosampler at 10 °C. Mobile solution A was 0.1% formic acid in water, and mobile phase B was 0.1% formic acid in acetonitrile. The run flow was 0.4 mL/min with gradient 20% B for 4 min, increasing linearly to 60% B for 2 min, holding at 60% B for 0.5 min, changing to 0% B in 0.5 min, and holding at 0% for 1 min. The mass data were collected on a Waters SQD2 detector with an m/z range of 2–3071.98 at a range of 1000–2000 m/z. The mass data were collected on a Waters SQD2 detector with an m/z range of 2–3071.98 at a range of 900–1500 m/z for ERK2 and 1000–2000 m/z for M^pro^. The desolvation temperature was 500 °C with a flow rate of 1000 L/h. The voltages used were 0.69 kV for the capillary and 46 V for the cone. Raw data were processed using openLYNX and deconvoluted using MaxEnt (20 - 60 kDa window, 1 Da/channel resolution).

### Kinase activity assays

Biochemical Kinase inhibition assays were carried out at Nanosyn, Santa Clara. Test compounds were diluted in 100% DMSO using 3-fold dilution steps. Final compound concentration in assay ranged from 10 μM to 0.0565 nM. Compounds were tested in a single well for each dilution, and the final concentration of DMSO in all assays was kept at 1%. Reference compound, Staurosporine, was tested in an identical manner. Compounds were preincubated in 25C for 2 hours before the measurements, and the kinase reactions were then performed for an additional 3 hours. For ERK2, the kinase concentration was 0.25-0.35 nM, the ATP concentration was 25 μM. For MELK, the kinase concentration was 0.06 nM, the ATP concentration was 30 μM. For VEGFR2, the kinase concentration was 0.25 nM, the ATP concentration was 80 μM. For GSK3B, the kinase concentration was 0.09 nM, the ATP concentration was 10 μM. For FGFR4, the kinase concentration was 0.17 nM, the ATP concentration was 250 μM.

### Biochemical M^pro^ inhibition assay

Compounds were seeded into assay-ready plates (Greiner 384 low volume 784900) using an Echo 555 acoustic dispenser, and DMSO was back-filled for a uniform concentration in assay plates (maximum 1%). Reagents for M^pro^ assay were dispensed into the assay plate in 10 μl volumes for a final of 20 μl. Final reaction concentrations were 20 mM HEPES pH=7.3, 1mM TCEP, 50 mM NaCl, 0.01% Tween-20, 10% glycerol, 5 nM M^pro^, 375 nM fluorogenic peptide substrate ([5-FAM]-AVLQSGFR-[Lys(Dabcyl)]-K-amide). M^pro^ was pre-incubated for 15 minutes at room temperature with compound before addition of substrate. Protease reaction was measured continuously in a BMG Pherastar FS with a 480/520 ex/em filter set. Data was mapped and normalized in Genedata Screener.

### M^pro^ Crystallography

M^pro^ protein was expressed and purified as discussed previously^47^. Apo M^pro^ crystals were grown using the sitting drop vapour diffusion method at 20 °C by adding 150 nl of protein (5 mg/ml in 20 mM Hepes pH 7.5, 50 mM NaCl) to 300 nl of crystallisation solution (11% PEG 4K, 6% DMSO, 0.1M MES pH 6.7) and 50 nl of seed stock prepared from initial crystal hits. 55 nl of a 100 mM compound stock solution in DMSO was added directly to the crystallisation drops using an ECHO liquid handler (final concentration 10% DMSO) and drops were incubated for approximately 1 hour prior to mounting and flash freezing in liquid nitrogen. Data were collected at Diamond Light Source on beamline I04-1 at 100K and processed using XDS^68^ and either xia2^69^, autoPROC^70^ or DIALS^71^. Further analysis was performed with XChemExplorer^72^: electron density maps were generated with Dimple^73^; ligand-binding events were identified using PanDDA^74^ (both the released version 0.2 and the pre-release development version (https://github.com/ConorFWild/pandda)); ligands were modelled into PanDDA-calculated event maps using Coot^75^; restraints were calculated with GRADE^76^; and structures were refined with BUSTER^77^. Coordinates, structure factors and PanDDA event maps for all data sets are deposited in the Protein Data Bank under PDB IDs 5RGT, 5RH5, 5RH6, 5RH7, 5RH9, 5RL0, 5RL1, 5RL2, 5RL3, 5RL4 and 5RL5. Data collection and refinement statistics are summarised in Supplementary Table 2. The ground-state structure and all corresponding datasets are deposited under PDB ID 5R8T.

## Supporting information

List of the Moonshot Consortium members

Supplementary Chemical Information

## Acknowledgments

We thank Prof. Oded Livnah for generously providing us recombinant ERK2. N.L. is the incumbent of the Alan and Laraine Fischer Career Development Chair. N.L. would like to acknowledge funding from the Israel Science Foundation (grants no. 2462/19 and 3824/19), The Israel Cancer Research Fund, the Israeli Ministry of Science Technology (grant no. 3-14763), the Moross Integrated Cancer Center and the Barry Sherman institute for Medicinal Chemistry. This research was supported by Nelson P. Sirotsky. N.L. is also supported by the Helen and Martin Kimmel Center for Molecular Design, Joel and Mady Dukler Fund for Cancer Research, the Estate of Emile Mimran and Virgin JustGiving, and the George Schwartzman Fund. D.Z. was funded in part by the pearlman student-initiated research award. The SGC is a registered charity (number 1097737) that receives funds from AbbVie, Bayer Pharma AG, Boehringer Ingelheim, Canada Foundation for Innovation, Eshelman Institute for Innovation, Genome Canada, Innovative Medicines Initiative (EU/EFPIA) [ULTRA-DD grant no. 115766], Janssen, Merck KGaA Darmstadt Germany, MSD, Novartis Pharma AG, Ontario Ministry of Economic Development and Innovation, Pfizer, São Paulo Research Foundation-FAPESP, Takeda, and Wellcome [106169/ZZ14/Z].

