## Supplementary Material for "An automatic pipeline for the design of irreversible derivatives identifies a potent SARS-CoV-2 M^pro^ inhibitor"



### Supplementary Figures

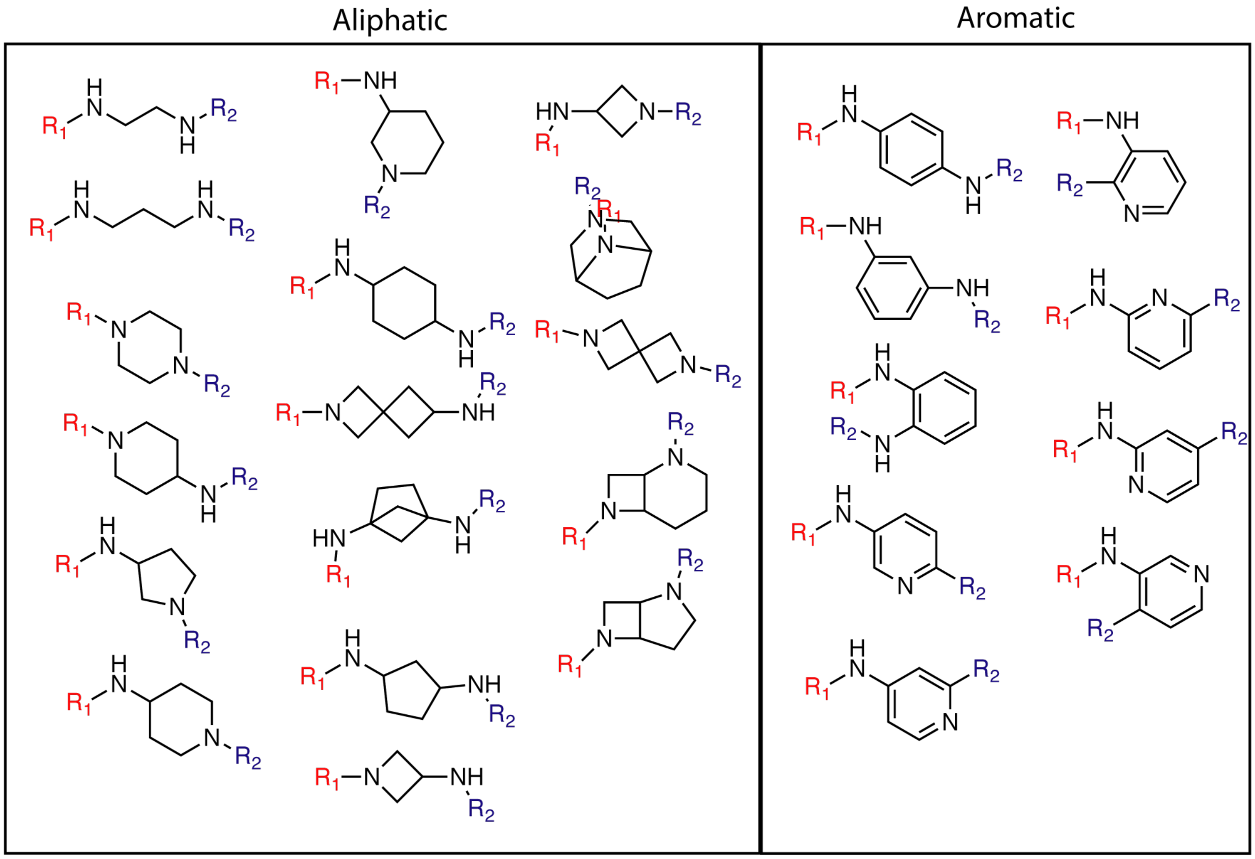


**Supplementary Figure 1. Di-amine linkers for installation of electrophiles.**
A list of di-amine linkers used in this manuscript to diversify the installation of electrophile unto reversible ligands. These were collected from the literature and were reported in the synthesis of covalent inhibitors [53–56]. R1 - Electrophile; R2 - Fragment from the reversible binder. See also: WO201821765.

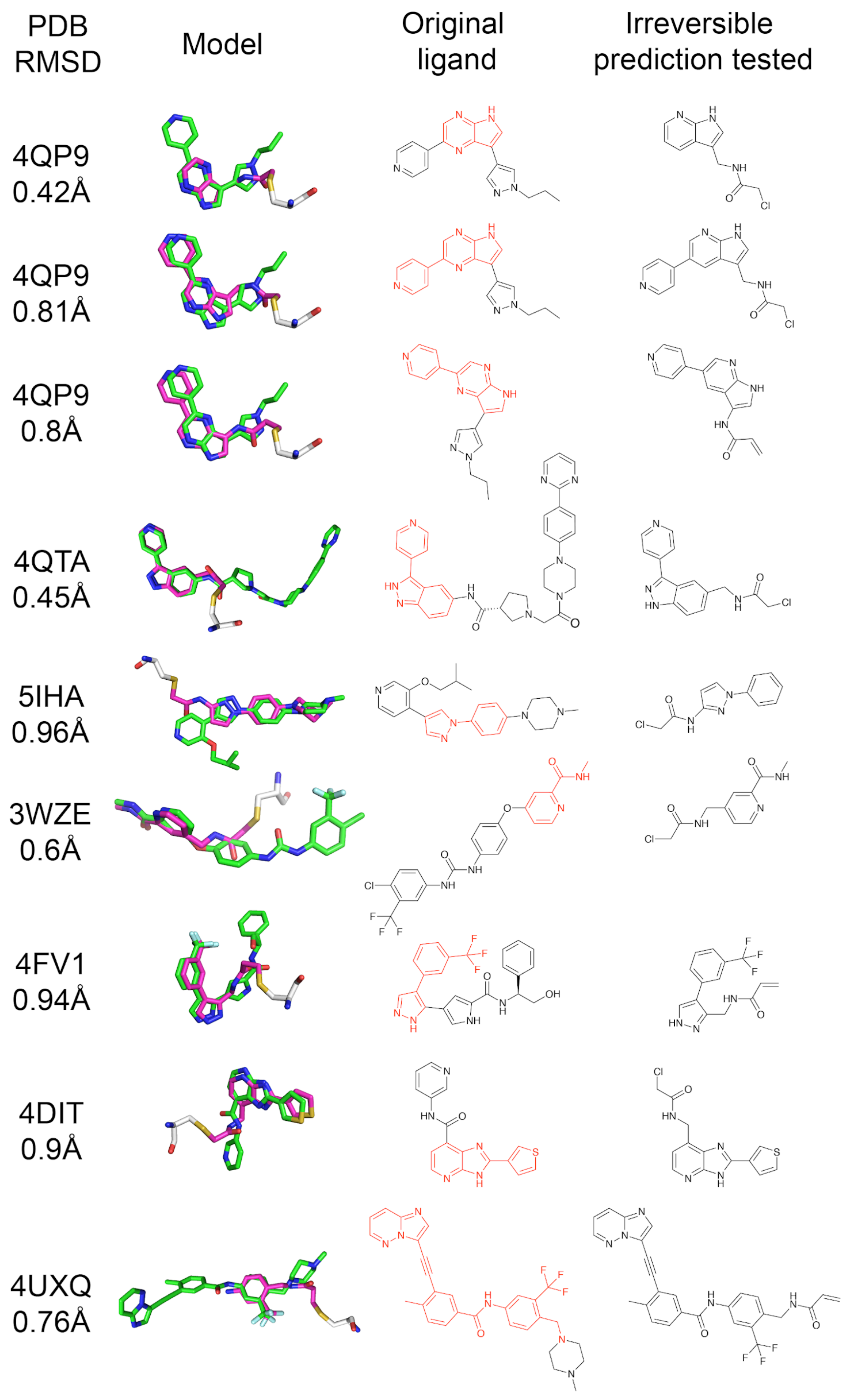


**Supplementary Figure 2. Prospective prediction of kinase covalent inhibitors.**

For each of the nine prospective covalent kinase inhibitors we made and synthesized we show: the original PDB on which it was based including MCS RMSD. A structural alignment of the covalentizer candidate (magenta) and the original reversible ligand (green). The 2D structure of the original reversible ligand (red indicates the substructure the prediction was based on) and the 2D structure of the covalent candidate.

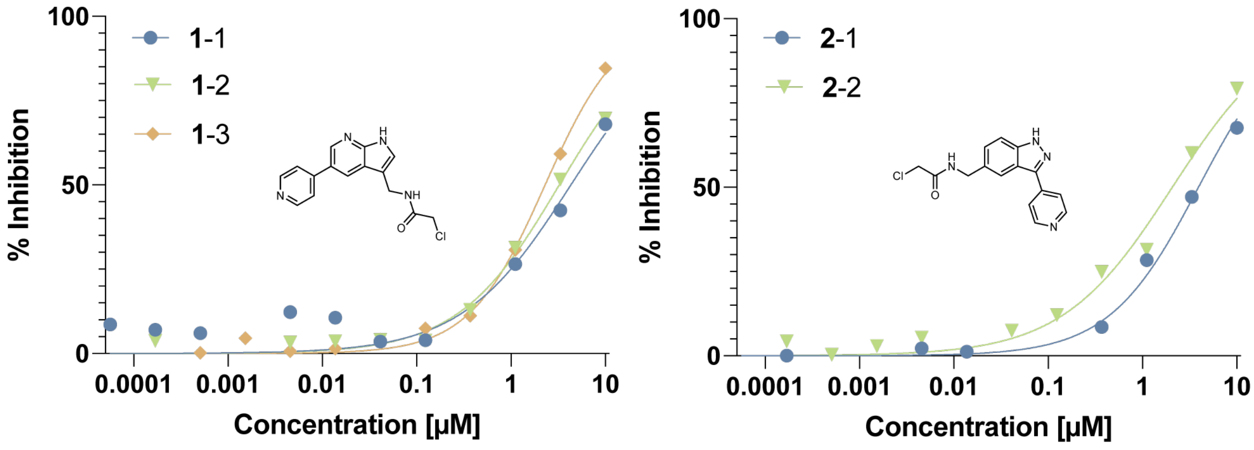


**Supplementary Figure 3. Additional $IC_{50}$ curves for two of the kinase inhibitors.**

Full $IC_{50}$ curves for ERK2 inhibition, for two of the nine kinase inhibitors reported in the manuscript. For **1**, the $IC_{50}$'s of the curves calculated from the data were 4.31 µM (**1**-1), 3.25 µM (**1**-2) and 2.3 µM (**1**-3) with an average of 3.29 µM. The assays were done by NanoSyn, Santa Clara. For **2**, the $IC_{50}$'s were 3.93 µM (**2**-1) and 2.07 µM (**2**-2), with an average of 3 µM.

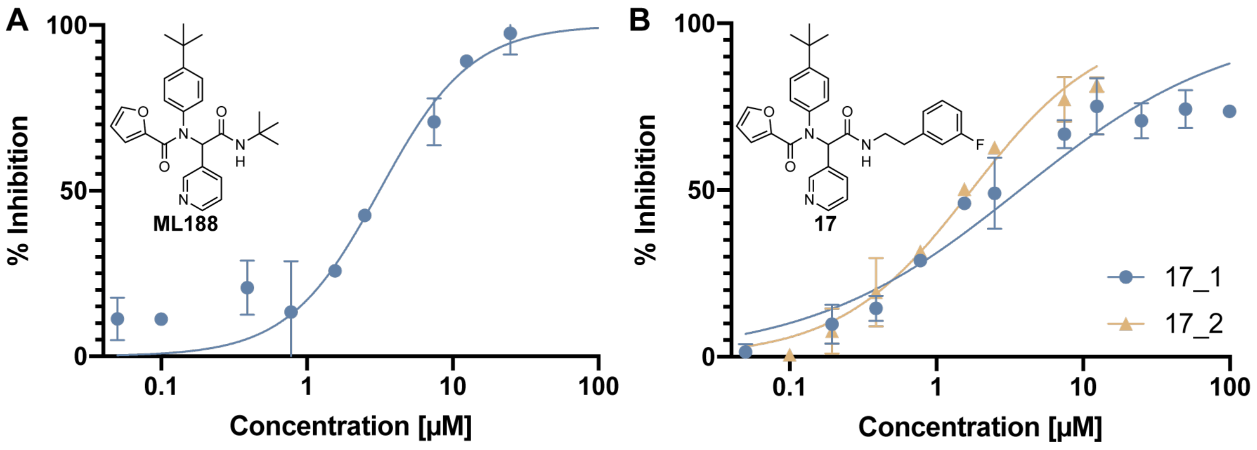


**Supplementary Figure 4. $IC_{50}$s of the reversible UGI compounds.**

**A.** Racemic **ML188** with an $IC_{50}$ of 3.14 µM. **B.** Compound **17** with an $IC_{50}$ of 3.71 µM (**17**-1) and 1.73 µM (**17**-2) with an average of 2.72 µM.

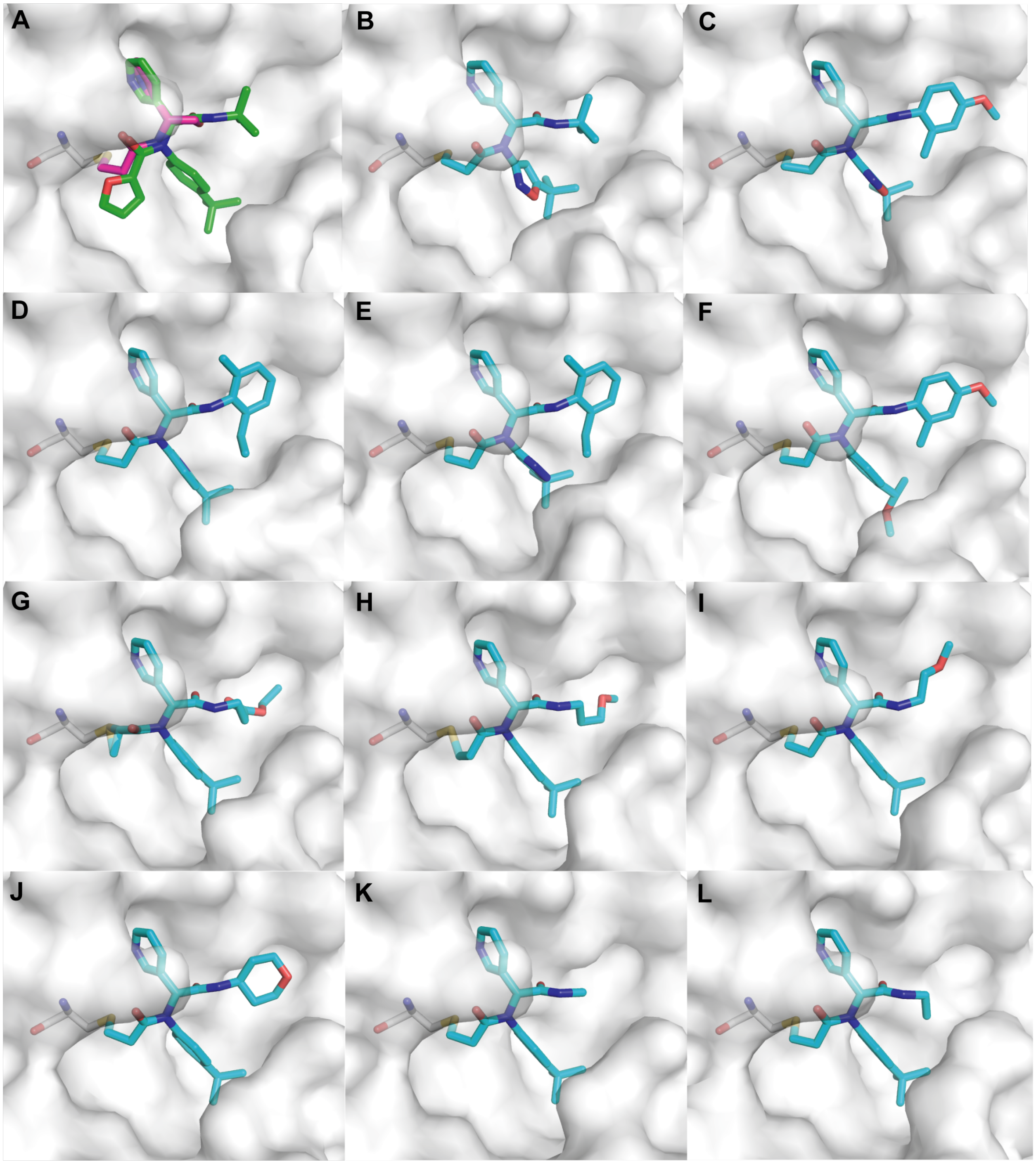


**Supplementary Figure 5. Crystal structures for analogs based on covalentizer predictions.** These are the crystal structures of 11 analogs of **10**. A. The covalentizer result, based on [40] (PDB: 3V3M). In green is the reversible ML188. In magenta is the covalentizer prediction. B. PDB: 5RGT. C. PDB: 5RH5. D. PDB: 5RH6. E. PDB: 5RH7. F. PDB: 5RH9. G. PDB: 5RL0. H. PDB: 5RL1. I. PDB: 5RL2. J. PDB: 5RL3. K. PDB: 5RL4. L. PDB: 5RL5.

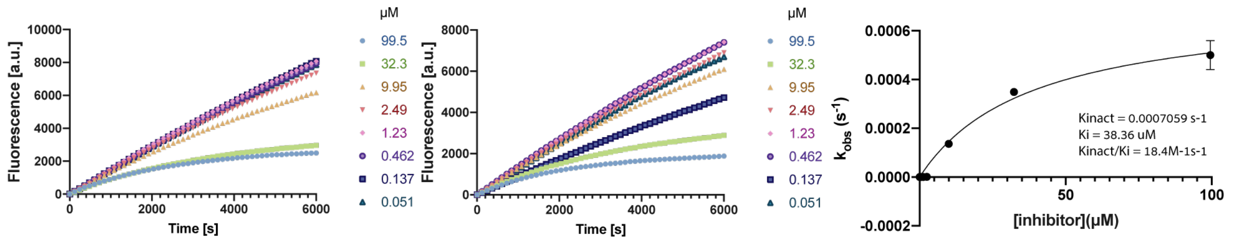


**Supplementary Figure 6. $K_i$/$K_{inact}$ of compound 11.**

Two repetitions (left and middle) of the fluorescence inhibition assay in a kinetic mode (no pre-incubation) with a range of inhibitor concentrations. Curves were fitted to this rate equation to calculate $k_{obs}$: Y=(va*X)+(vb-va)*(1-EXP(-$k_{obs}$*X))/($k_{obs}$).The right panel plots the $k_{obs}$ as a function of inhibitor concentration. Each point is an average of the two values from the two repetitions on the left. $K_i$ and $K_{inact}$ were extracted by fitting Y=$K_{inact}$*X/($K_i$+X).

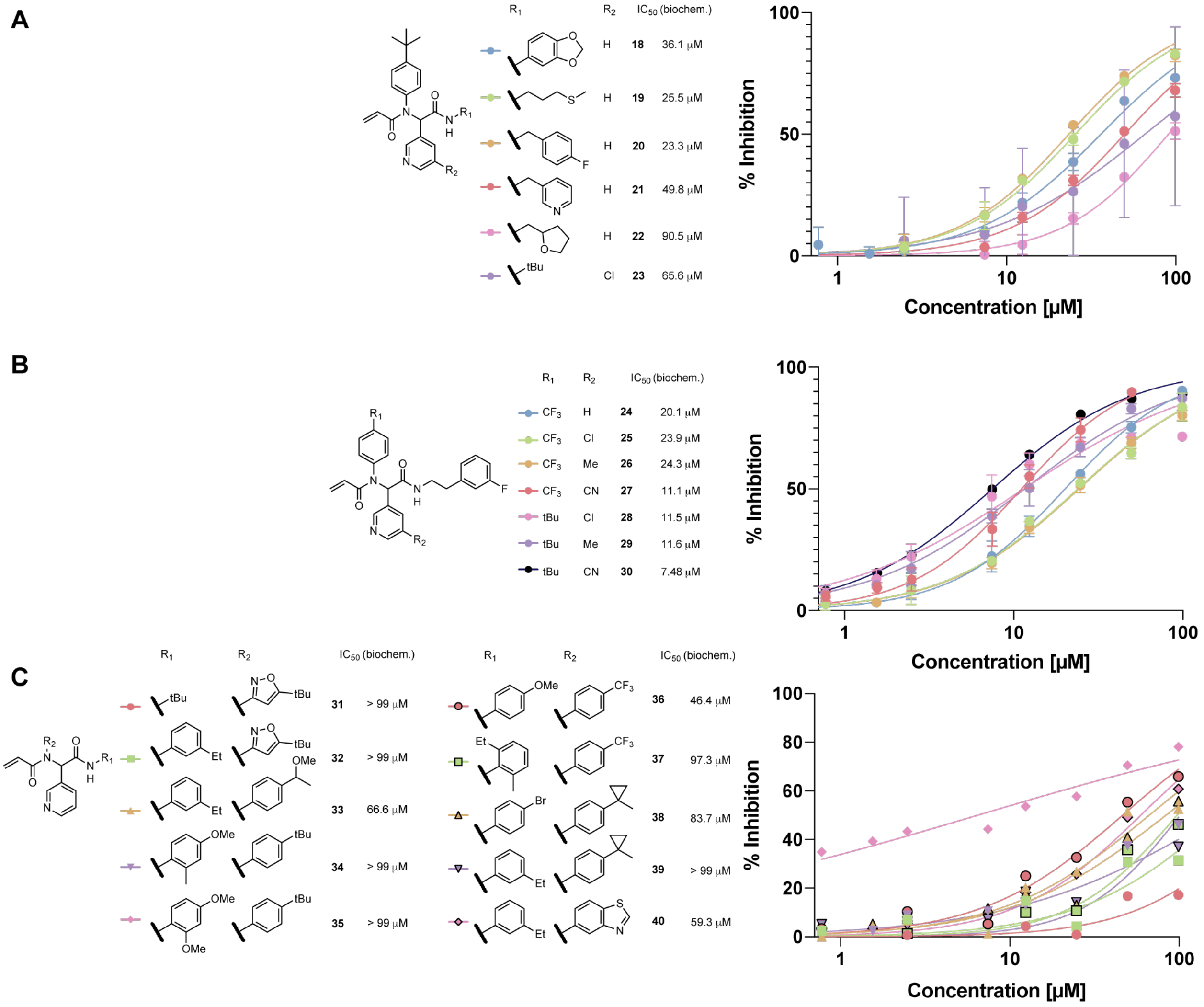


**Supplementary Figure 7. SAR analysis of additional M$^{pro}$ covalent inhibitors.**

**A.** Biochemical $IC_{50}$'s and their associated curves for early inhibitors probing the S3 pocket. **B.** Biochemical $IC_{50}$'s and their associated curves for inhibitors with a 3'-fluorophenethylamide motif optimized for the S3 pocket combined with independently optimized substituents for the S1 and S2 sub-pockets showing non-synergistic effects. **C.** Biochemical $IC_{50}$'s and their associated curves for early combinatorial probing of the S1 and S3 sub-pockets.

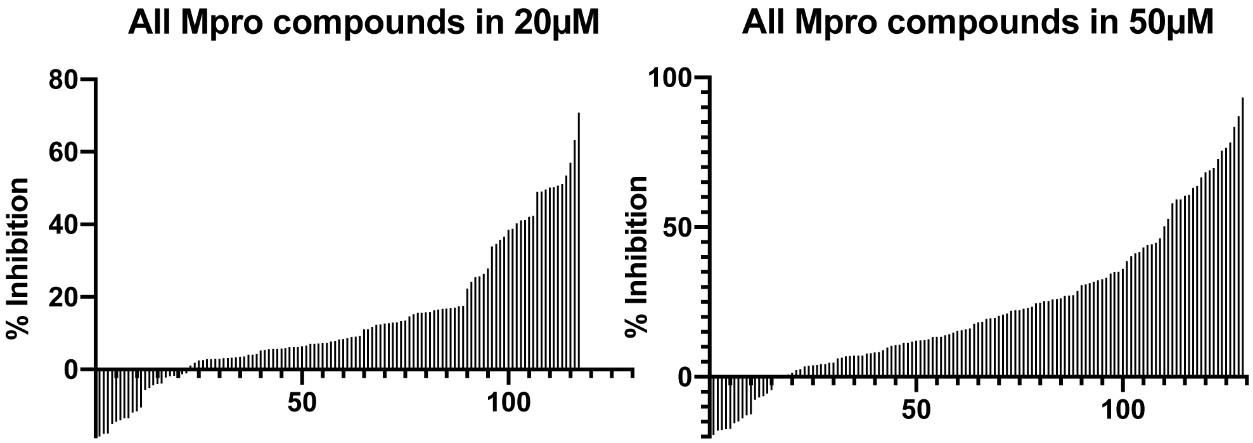


**Supplementary Figure 8. Single point inhibition by 130 covalent M$^{pro}$ inhibitors (20/50 µM).** % inhibition at two single doses: 20 µM (left) and 50 µM (right), for the majority of the 130 analogs of **10** which we made and tested. There is a large variability in inhibition over this large set of analogs, starting from inactive compounds, up to compounds that inhibit 60-80% in 20 µM. The date is available in Supp. Dataset 1.

### Supplementary Tables

| Warhead | PDB ID | Best RMSD (Å)[a] | Warhead | PDB ID | Best RMSD (Å) |
|---|---|---|---|---|---|
| Acrylamides | 2HWO | 1.91* | Acrylamides | 5TOZ | 0.69 |
| | 4HCU | 1.83 | | 5TTS | 1.05 |
| | 4HCV | 0.81 | | 5TTU | 0.91 |
| | 4QPS | 1.59* | | 5TTV | 2.93 |
| | 4XCU | 1.34 | | 5UG8 | 1.38 |
| | 4ZZO | 0.35 | | 5UGC | 2.08 |
| | 5GNK | 1.09 | Substituted Acrylamides | 3T9T | 3.53 |
| | 5HG5 | 1.76 | | 3W2P | 3.45* |
| | 5HG7 | 2.8 | | 4I24 | 0.47 |
| | 5HG8 | 1.8 | | 4YHF | 6.66* |
| | 5HG9 | 2.23 | | 5P9M | 1.44* |
| | 5J7S | 2.2* | Chloro-acetamides | 5E7R | 3.48* |
| | 5J87 | 1.49 | | 5L6P | 0.5 |
| | 5J9Z | 0.97 | | 5MJB | 1.6* |
| | 5JK3 | 1.98* | Sulfonamides | 3SVV | 3.67* |
| | 5LCK | 0.81 | | 4ZZM | 4.82* |
| | 5O8V | 1.99 | | | |
| | 5P9J | 0.8 | | | |
| | 5P9K | 2.36 | | | |

**Supplementary Table 1. Covalent kinase inhibitor benchmark,**

A kinase subset of a recently published covalent docking benchmark [30].

[a] RMSD after removing the electrophile from the covalent inhibitor (leaving only a free amine), and re-docking using covalentizer.

* In these cases the fragment does not recapitulate the covalent warhead attachment.

| | Mpro-x2694 | Mpro-x2703 | Mpro-x2705 | Mpro-x2776 | Mpro-x3115 | Mpro-x3117 | Mpro-x3124 | Mpro-x3113 | Mpro-x3110 | Mpro-x3359 | Mpro-x2540 |
|---|---|---|---|---|---|---|---|---|---|---|---|
| Wavelength | 0.9126 ≈ | 0.9126 ≈ | 0.9126 ≈ | 0.9126 ≈ | 0.9126 ≈ | 0.9126≈ | 0.9126 ≈ | 0.9126 ≈ | 0.9126 ≈ | 0.9126 ≈ | 0.9126 ≈ |
| Resolution range | 54.76 - 1.72 (1.781 - 1.72) | 47.84 - 1.6 (1.657 - 1.6) | 47.69 - 1.71 (1.771 - 1.71) | 47.58 - 1.91 (1.978 - 1.91) | 54.94 - 1.48 (1.533 - 1.48) | 55.08 - 1.51 (1.564 - 1.51) | 47.44 - 1.53 (1.585 - 1.53) | 47.49 - 1.65 (1.709 - 1.65) | 54.85 - 1.69 (1.75 - 1.69) | 55.52 - 1.58 (1.636 - 1.58) | 47.55 - 2.218 (2.298 - 2.218) |
| Space group | C 1 2 1 | C 1 2 1 | C 1 2 1 | C 1 2 1 | C 1 2 1 | C 1 2 1 | C 1 2 1 | C 1 2 1 | C 1 2 1 | C 1 2 1 | C 1 2 1 |
| Unit cell | 112.37 52.92 44.41 90 102.95 90 | 112.79 8 53.132 44.453 90 102.82 3 90 | 112.60 3 52.957 44.456 90 102.99 3 90 | 112.74 2 52.785 44.39 90 103.02 2 90 | 112.78 9 52.691 44.462 90 103.05 1 90 | 113.059 52.832 44.547 90 103.017 90 | 112.25 1 52.644 44.3 90 102.83 6 90 | 112.36 9 52.693 44.357 90 102.87 3 90 | 112.56 1 52.771 44.375 90 102.96 5 90 | 113.82 9 53.253 44.449 90 102.71 90 | 112.39 6 52.781 44.792 90 103.05 1 90 |
| Total reflections | 99064 (8800) | 114118 (9166) | 95332 (8271) | 69990 (7120) | 131594 (8738) | 127200 (8677) | 12286 9 (8393) | 102917 (8472) | 98472 (8314) | 118260 (9243) | 48314 (4978) |
| Unique reflections | 27080 (2663) | 33779 (2805) | 27345 (2332) | 19770 (1771) | 40458 (3618) | 38593 (3475) | 37636 (3239) | 30135 (2822) | 28378 (2710) | 35454 (3119) | 12786 (1275) |
| Multiplicity | 3.7 (3.3) | 3.4 (2.8) | 3.5 (3.1) | 3.5 (3.6) | 3.3 (2.4) | 3.3 (2.5) | 3.3 (2.4) | 3.4 (2.9) | 3.5 (3.0) | 3.3 (2.7) | 3.8 (3.9) |
| Completeness (%) | 99.64 (98.81) | 97.67 (84.01) | 97.07 (84.77) | 97.89 (90.40) | 95.22 (85.63) | 97.90 (86.70) | 97.90 (85.15) | 97.93 (92.85) | 99.21 (96.41) | 98.23 (89.01) | 99.70 (99.61) |
| Mean I/sigma(I) | 8.28 (0.71) | 8.62 (0.43) | 6.97 (0.44) | 4.09 (0.42) | 9.80 (0.56) | 10.25 (0.66) | 8.59 (0.46) | 7.19 (0.46) | 9.29 (0.62) | 8.72 (0.38) | 5.23 (1.04) |
| Wilson B-factor | 26.6 | 24.02 | 24.03 | 27.13 | 18.24 | 19.81 | 19.65 | 21.74 | 23.29 | 24.11 | 34.75 |
| R-merge | 0.0882 6 (1.653) | 0.0785 6 (1.28) | 0.1043 (1.2) | 0.212 (2.02) | 0.0721 8 (1.019) | 0.07103 (0.8922 ) | 0.0834 5 (1.259) | 0.1094 (1.455) | 0.0943 9 (1.348) | 0.0733 4 (1.483) | 0.1934 (1.384) |

| | | | | | | | | | | | |
|---|---|---|---|---|---|---|---|---|---|---|---|
| R-meas | 0.1036 (1.977) | 0.09315 (1.582) | 0.1233 (1.452) | 0.2497 (2.362) | 0.0858 (1.292) | 0.08431 (1.119) | 0.09888 (1.554) | 0.1294 (1.782) | 0.1113 (1.628) | 0.08701 (1.846) | 0.2254 (1.598) |
| R-pim | 0.05343 (1.066) | 0.04926 (0.9158) | 0.0648 (0.802) | 0.1299 (1.205) | 0.04553 (0.7783) | 0.04462 (0.6609) | 0.05216 (0.895) | 0.06781 (1.006) | 0.058 (0.9001) | 0.04607 (1.08) | 0.1145 (0.7922) |
| CC1/2 | 0.998 (0.373) | 0.996 (0.357) | 0.995 (0.33) | 0.971 (0.317) | 0.998 (0.357) | 0.998 (0.48) | 0.996 (0.367) | 0.997 (0.401) | 0.997 (0.362) | 0.998 (0.363) | 0.989 (0.399) |
| CC* | 1 (0.737) | 0.999 (0.725) | 0.999 (0.705) | 0.993 (0.694) | 0.999 (0.725) | 0.999 (0.805) | 0.999 (0.732) | 0.999 (0.757) | 0.999 (0.729) | 0.999 (0.73) | 0.997 (0.755) |
| Reflections used in refinement | 27035 (2658) | 33151 (2805) | 26877 (2332) | 19442 (1770) | 40423 (3617) | 39436 (3475) | 37333 (3239) | 29916 (2820) | 28313 (2710) | 35016 (3117) | 12773 (1275) |
| Reflections used for R-free | 1335 (120) | 1645 (143) | 1327 (98) | 1000 (84) | 1956 (165) | 1922 (146) | 1828 (150) | 1469 (135) | 1397 (125) | 1727 (146) | 653 (56) |
| R-work | 0.1937 (0.3912) | 0.1975 (0.3639) | 0.2016 (0.3734) | 0.2004 (0.3579) | 0.1853 (0.3495) | 0.1847 (0.3488) | 0.1956 (0.3830) | 0.2008 (0.4120) | 0.1873 (0.3542) | 0.1994 (0.3934) | 0.1826 (0.2572) |
| R-free | 0.2325 (0.4356) | 0.2330 (0.3844) | 0.2404 (0.4345) | 0.2572 (0.3779) | 0.2034 (0.3316) | 0.2123 (0.3506) | 0.2210 (0.3532) | 0.2320 (0.3929) | 0.2247 (0.3931) | 0.2352 (0.4139) | 0.2582 (0.3789) |
| CC(work) | 0.959 (0.614) | 0.961 (0.631) | 0.961 (0.605) | 0.957 (0.631) | 0.965 (0.627) | 0.959 (0.620) | 0.962 (0.638) | 0.961 (0.668) | 0.964 (0.648) | 0.964 (0.593) | 0.951 (0.700) |
| CC(free) | 0.957 (0.558) | 0.967 (0.520) | 0.961 (0.438) | 0.956 (0.368) | 0.973 (0.667) | 0.958 (0.698) | 0.969 (0.706) | 0.971 (0.643) | 0.961 (0.577) | 0.960 (0.573) | 0.949 (0.382) |
| Number of non-hydrogen atoms | 2675 | 2707 | 2678 | 2649 | 2734 | 2742 | 2713 | 2667 | 2708 | 2709 | 2571 |
| macromolecules | 2360 | 2360 | 2363 | 2360 | 2360 | 2360 | 2360 | 2360 | 2360 | 2355 | 2360 |
| ligands | 49 | 49 | 49 | 50 | 45 | 47 | 42 | 46 | 52 | 43 | 44 |
| solvent | 266 | 298 | 266 | 239 | 329 | 335 | 311 | 261 | 296 | 311 | 167 |

| | | | | | | | | | | | |
|---|---|---|---|---|---|---|---|---|---|---|---|
| Protein residues | 304 | 304 | 304 | 304 | 304 | 304 | 304 | 304 | 304 | 304 | 304 |
| RMS(bonds) | 0.012 | 0.012 | 0.012 | 0.012 | 0.011 | 0.011 | 0.011 | 0.012 | 0.29 | 0.327 | 0.012 |
| RMS(angles) | 1.59 | 1.56 | 1.58 | 1.6 | 1.57 | 1.57 | 1.59 | 1.58 | 7.27 | 6.4 | 1.61 |
| Ramachandran favored (%) | 98.01 | 97.35 | 97.35 | 96.36 | 97.35 | 98.68 | 97.68 | 97.35 | 97.35 | 98.01 | 96.36 |
| Ramachandran allowed (%) | 1.66 | 2.32 | 2.32 | 2.98 | 2.32 | 0.99 | 1.99 | 1.99 | 2.32 | 1.66 | 3.31 |
| Ramachandran outliers (%) | 0.33 | 0.33 | 0.33 | 0.66 | 0.33 | 0.33 | 0.33 | 0.66 | 0.33 | 0.33 | 0.33 |
| Rotamer outliers (%) | 1.14 | 0.76 | 0.38 | 1.52 | 1.9 | 0.38 | 1.9 | 2.28 | 1.14 | 0 | 2.66 |
| Clashscore | 2.32 | 1.9 | 2.52 | 3.79 | 2.53 | 3.16 | 2.53 | 2.95 | 2.31 | 1.69 | 3.58 |
| Average B-factor | 33.74 | 29.74 | 30.06 | 35.63 | 23.99 | 25.3 | 26.55 | 29.65 | 29.33 | 30.46 | 40.12 |
| macromolecules | 32.32 | 28.2 | 28.87 | 34.93 | 22.23 | 23.49 | 24.84 | 28.36 | 27.66 | 28.89 | 39.6 |
| ligands | 48.55 | 39.12 | 40.42 | 44.86 | 31.59 | 35.84 | 33.02 | 38.78 | 44.41 | 43.91 | 65.65 |
| solvent | 43.64 | 40.42 | 38.75 | 40.56 | 35.62 | 36.58 | 38.64 | 39.71 | 40.01 | 40.5 | 40.8 |

**Supplementary Table 2. Crystallographic statistics table.**
