## Supplementary Chemical Information for "An automatic pipeline for the design of irreversible derivatives identifies a potent SARS-CoV-2 M^pro^ inhibitor"

### Organic Synthesis Methods

Unless otherwise noted, all reactions were performed in air with analytical grade solvents. All reagents and solvents used for the synthesis were used as acquired without further purification. Compounds **3, 4, 7, 9, 11, 12, 13, 14** and all other Ugi products were acquired from Enamine (custom synthesis). Flash chromatography was performed using a CombiFlash® System (Teledyne Isco, USA) with RediSep Rf Normal-phase Flash Columns. Reversed phase preparative HPLC was carried out using a Waters Prep 2545 Preparative Chromatography System, with UV/Vis detector 2489 at variable wavelengths (specified below), using XBridge® Prep C18 10µm 10x250 mm Column (PN: 186003891, SN:161I3608512502), 10 mL/min flow rate. Reaction progress and compounds' purity was monitored by Waters UPLC-MS system: Acquity UPLC® H class with PDA detector, and using Acquity UPLC® BEH C18 1.7 µm 2.1x50 mm Column (PN:186002350, SN 02703533825836). MS-system: Waters, SQ detector 2. TLC reaction progress analysis was carried out with aluminium backed $SiO_2$ plates with F254 indicator, and compounds were visualized either under UV light or with the typical staining methods.

$^{1}H$- and $^{13}C$-NMR spectra were recorded on a Bruker Avance III -300 MHz, 400 MHz spectrometer, equipped with a QNP probe. Chemical shifts are reported in ppm on the δ scale down field from TMS and are calibrated according to the deuterated solvents. All J values are given in Hertz. High resolution electron-spray mass spectrometry (HR-MS) of final compounds was performed by the Department of Chemical Research Support, Weizmann Institute.

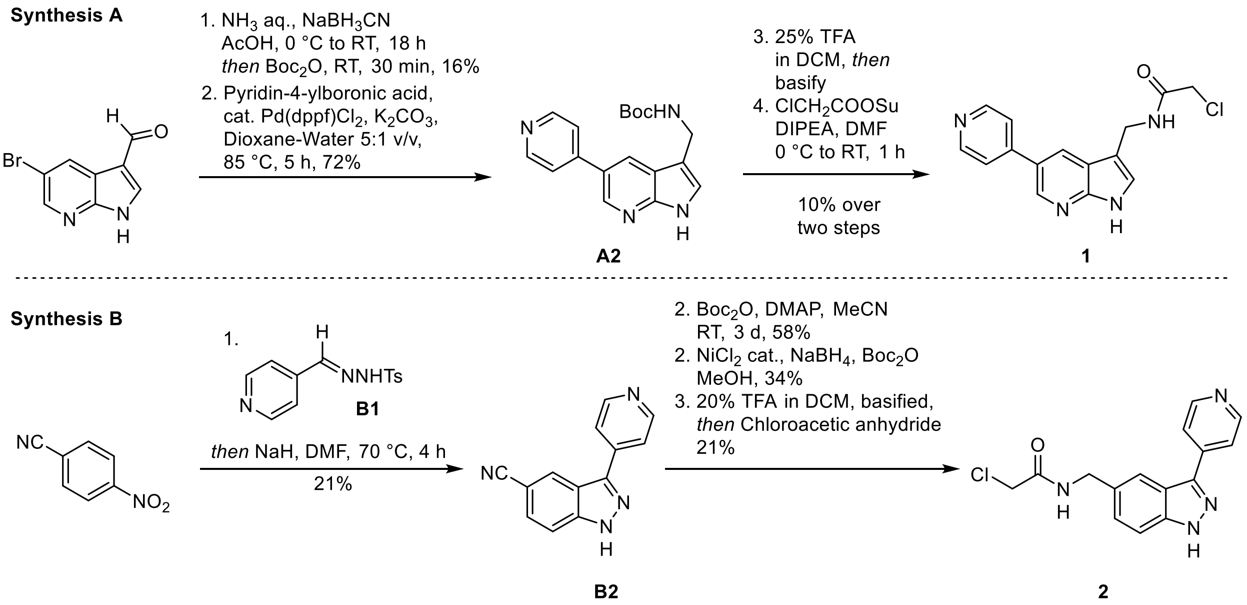


**Supplementary Scheme 1. Synthetic routes towards active irreversible kinase inhibitors designed by the Covalentizer computational pipeline. Boc: tert-butyloxycarbonyl, DIPEA: Diisopropylethylamine, dppf: 1,1'-Bis(diphenylphosphino)ferrocene, Su: Succinimide, TFA: Trifluoroacetic acid.**

**Synthesis A:** *N*-Boc-(5-bromo-1*H*-pyrrolo[2,3-b]pyridin-3-yl)methanamine (**A1**)

A 25 mL RBF with stir bar was charged with 5-bromo-1H-pyrrolo[2,3-b]pyridine-3-carbaldehyde (112.5 mg, 500 µmol, 1.00 equiv.) and glacial acetic acid (7 mL). The stirred solution was then cooled to 0 °C using an ice bath, followed by the slow addition of 25% aq. ammonia (4.18 mL, 54 mmol, 108 equiv.) and finally sodium cyanoborohydride (94.3 mg, 1.50 mmol, 3.00 equiv.) at the same temperature. The ice bath was allowed to warm overnight and the reaction mixture, containing a suspension, thoroughly stirred. After 16 h, the suspension had disappeared and the clear solution was basified to pH 12 using 4 M aq. NaOH at 0 °C. The aqueous layer was extracted with *i*PrOH:$CHCl_3$ 1:3 (v/v, 3 x 20 mL). The combined organic phases were dried over $MgSO_4$, filtered and the resultant solution concentrated (ca. 5 mL) using a rotary evaporator. The concentrate was transferred to a 20 mL vial, followed by the addition of $Boc_2O$ (600 µL, ca. 2.5 mmol, 5 equiv. assuming complete conversion to the free amine). After 30 min, LC-MS analysis of the reaction mixture revealed a mixture of the product with some byproducts, including the corresponding free primary alcohol, 3-methyl-5-bromo-7-azaindole as well as the *N*-Boc protected secondary amine dimer. The solvent and excess reagent were removed by evaporation *in vacuo*. The crude product was purified by flash chromatography of a wet-loaded DCM extract of the crude solid (stationary phase: $SiO_2$, mobile phase: gradient from 0 to 8% MeOH in $CHCl_3$) to give the product as a fine white powder (18 mg, 80 µmol, 16%).

LC-MS: m/z 327 corresponding to $[M+H]^+$, m/z 329 corresponding to $[M+H]^+$ for the heavier bromine isotope.

**Synthesis A:** *N*-Boc-(5-pyrid-4-yl-1*H*-pyrrolo[2,3-b]pyridin-3-yl)methanamine (**A2**)

The heteroaryl bromide **A1** (18 mg, 55.2 µmol, 1 equiv.) was charged into a 2 dram vial containing a stir bar, pyrid-4-yl-boronic acid (8.2 mg, 66.2 µmol, 1.2 equiv.) and solid $K_2CO_3$ (22.9 mg, 165.55 µmol, 3.0 equiv.). The solids were dissolved in 1,4-dioxane (2 mL) and water (0.3 mL). The resultant stirred solution was thoroughly sparged with Ar gas for 15 min, followed by the addition of Pd(dppf)$Cl_2$ (4 mg, 5.52 µmol, 10 mol%) under an Ar stream. The solution was further sparged with Ar for 1 min. The vial was sealed, the solution stirred and heated to 85 °C using an oil bath. The initial yellow colored solution changed to a deep dark-red color. After 5 h, TLC and LC-MS analysis indicated complete conversion to the product. The reaction mixture was cooled to RT and diluted with water (1 mL) and EtOAc (2 mL). The phases were separated and the aq. layer extracted with EtOAc (2 x 1 mL). The combined organic phases were dried over $Na_2SO_4$ and then filtered in a Pasteur pipette filled with a plug of cotton. The plug was washed with a small amount of MeOH. The filtrate was concentrated to dryness by rotary evaporation, giving a brown solid residue as the product (13 mg, 40.1 µmol, 73%) which was directly employed in the next step without further purification.

LC-MS: m/z 325 corresponding to $[M+H]^+$.

**Synthesis A:** *N*-Chloroacetamido-(5-pyridin-4-yl-1*H*-pyrrolo[2,3-b]pyridin-3-yl)methanamine **(1)**

*N*-Boc-amine **A2** (22 mg, 67.44 µmol, 1 equiv.) was suspended in DCM (1.5 mL) in a 2 dram vial under stirring, cooled to 0 °C using an ice bath, followed by the slow addition of TFA (0.5 mL) at the same temperature, which resulted in complete dissolution to give a cherry-red reaction mixture. After 10 min, the ice bath was removed and the reaction mixture allowed to stir at RT for 2 h. Excess solvent and TFA was removed by passing a stream of Ar over the reaction mixture. The residual oil was carefully treated with $NaHCO_3$ until basic and then extracted from the slurry using *i*PrOH:DCM 1:3 v/v (2 x 2 mL). The organic phase was dried over a small amount of $MgSO_4$ for 15 min, filtered through a cotton plug in a pasteur pipette and freed from solvent by rotary evaporation. The vial containing the dried free base was placed under an Ar atmosphere, followed by the dissolution of the oil in dry DMF. The dark solution was treated with DIPEA (12 µL, 67.44 µmol, 1.0 equiv.), cooled to 0 °C followed by the addition of chloroacetic acid NHS ester (13 mg, 80.9 µmol, 1.2 equiv) under a gentle Ar stream. After 1 h, the reaction was deemed complete by LC/MS and stopped by the addition of glacial AcOH (20 µL). Bulk DMF was removed by rotary evaporation under vacuum at 70 °C water bath temperature. The dark residue was partially dissolved in 50% aq. ACN (4 mL) and filtered through a 0.2 µm PVDF syringe tip filter. The filtrate was injected for purification by preparative RP-HPLC (C18 column, 5 to 95% MeCN in $H_2O$ w. 0.1% TFA over 40 min, 10 mL/min, $\lambda$ = 254 nm). After lyophilization of the target fractions, the product was obtained as a white solid (2.94 mg, 9.44 µmol, 14%).

$^1$H-NMR (400 MHz, DMSO-$d_6$) $\delta$ ppm 11.95 (s, 1H), 8.85 – 8.92 (m, 3H), 8.65 – 8.71 (m, 2H), 8.30 (d, *J* = 6.5 Hz, 2H), 7.55 (d, *J* = 2.1 Hz, 1H), 4.52 (d, *J* = 5.5 Hz, 2H), 4.10 (s, 2H). $^{13}$C-NMR (101 MHz, DMSO-$d_6$): $\delta$ ppm 166.3, 158.7, 158.4, 153.3, 150.2, 144.7, 143.0, 127.3, 126.9, 123.0, 119.3, 112.8, 43.2, 34.8. LC-MS: m/z 301 corresponding to $[M+H]^+$, m/z 303 corresponding to $[M+H]^+$ for the heavier chlorine isotope. HRMS: m/z, $C_{15}H_{13}ClN_4O$, required 301.0856 $[M+H]^+$, found 301.0860 $[M+H]^+$, 1.3 ppm deviation.

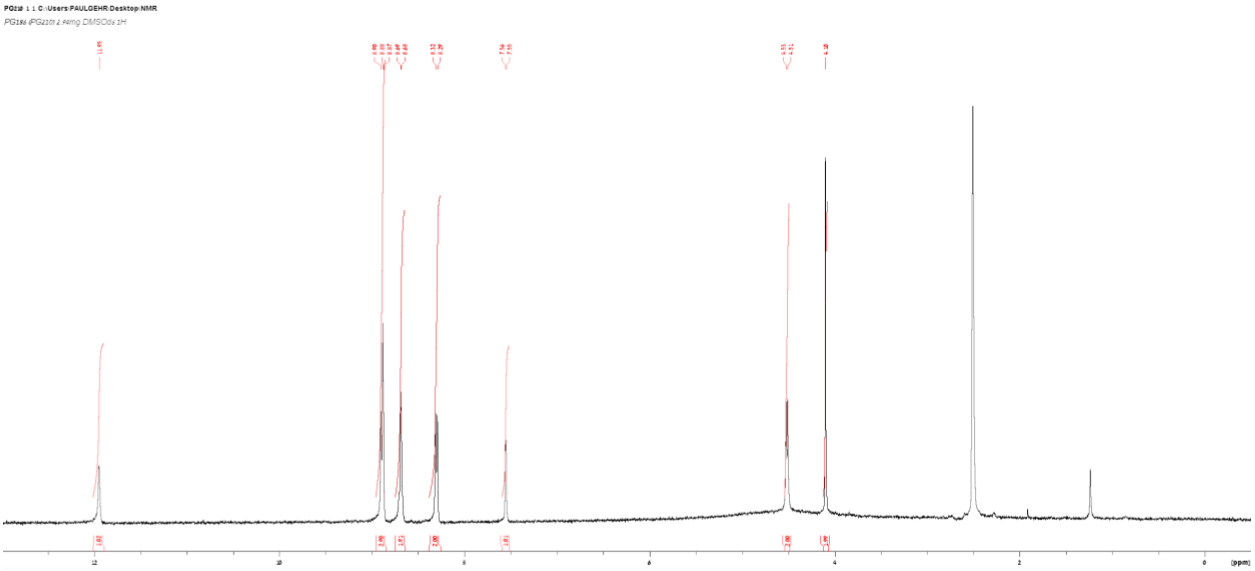

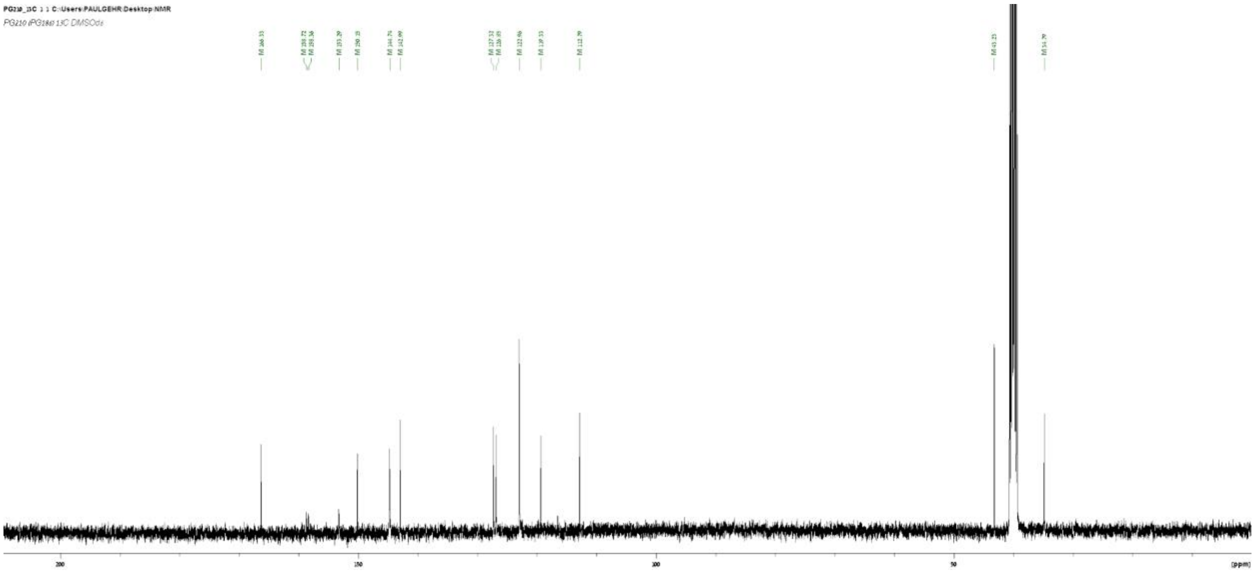

**Synthesis B:** 4-Methyl-N'-(pyridin-4-ylmethylene)benzenesulfonohydrazide (**B1**)

*N*-tolylhydrazine (1.98 g, 10.63 mmol, 1 equiv.) was dissolved in warm EtOH (8 mL), followed by the addition of isonicotinaldehyde (1 mL, 10.63 mmol, 1.0 equiv.) and the reaction mixture was cooled to 0 °C, reacted for 30 min and then vacuum filtered and washed with hexane and diethyl ether until the filter cake was nearly white. The yellow filtrate was discarded. The filter cake was dried to afford the product as an off-white solid (2.3 g, 7.76 mmol, 73%).

LC-MS: m/z 276 corresponding to $[M+H]^+$.

**Synthesis B:** 3-(Pyridin-4-yl)-1*H*-indazole-5-carbonitrile (**B2**)

The hydrazone **B1** (1.00 g, 3.63 mmol, 1 equiv.) and 4-nitrobenzonitrile (0.645 g, 4.36 mmol, 1.2 equiv.) were dissolved in dry DMF (24 mL) under Ar in a dried reaction vessel. NaH (522 mg of a 60% w/w oil dispersion, 13.08 mmol, 3.6 equiv.) was added to the mixture under a stream of Ar. The reaction vessel was stirred at 80 °C for 4 h under an Ar atmosphere, then cooled to RT, quenched with 1 M HCl until excess NaH was fully consumed. The reaction mixture was diluted with brine and then basified with 5% $NaHCO_3$. The solution was extracted with *i*PrOH:$CHCl_3$ 1:3 v/v (2 x 25 mL), then dried over $MgSO_4$, the volatiles removed by rotary evaporation. The crude product was thoroughly dried in vacuo, adsorbed on Celite by evaporation from a solution in EtOAc and MeOH, and then purified by flash chromatography (24 g $SiO_2$, 0 to 50% *i*PrOH in $CHCl_3$) to give the product as an orange solid (166 mg, 0.754 mmol, 21%).

LC-MS: m/z 221 corresponding to $[M+H]^+$.

**Synthesis B:** $N^1$-Boc-3-(Pyridin-4-yl)-1*H*-indazole-5-carbonitrile (**B3**)

**B2** (110 mg, 495.4 µmol, 1.0 equiv.) was dissolved in MeCN (5 mL), followed by addition of DMAP (60.5 mg, 495.4 µmol, 1.0 equiv.) and then $Boc_2O$ (171 µL, 743.1 µmol, 1.5 equiv.) and stirred for 3 d at RT. The crude reaction mixture was adsorbed on celite and purified by flash chromatography (4 g $SiO_2$, 0 to 50% EtOAc in Hexane) to give the product (93 mg, 0.288 µmol, 58%).

LC-MS: m/z 321 corresponding to $[M+H]^+$.

**Synthesis B:** $N^1$-Boc-3-(Pyridin-4-yl)-1*H*-indazole-5-carbonitrile (**B4**)

**B3** (30 mg, 94 µmol, 1.0 equiv.) was dissolved in MeOH, cooled to 0 °C, followed by addition of $NiCl_2 \bullet 6 H_2O$ (4.5 mg, 18.8 µmol, 20 mol%), Boc2O (43 µL, 187 µmol, 2.0 equiv.) and finally $NaBH_4$ (24.8 mg, 655.5 µmol, 7.0 equiv.) in several portions to avoid excessive bubbling. The reaction was allowed to warm to room temperature overnight. MeOH was removed by rotary evaporation and the residue partitioned between aq. sat. $NaHCO_3$ and EtOAc. The organic phase was dried by filtration through a pipette plug of $MgSO_4$, adsorbed on celite and purified by flash chromatography (4 g $SiO_2$, 0 – 100% EtOAc in Hexane) to give the product (7 mg, 16.5 µmol, 18%).

LC-MS: m/z 425 corresponding to $[M+H]^+$.

**Synthesis B:** *N*-((3-(Pyridin-4-yl)-1H-indazol-5-yl)methyl)chloroacetamide (**2**)

Intermediate **B4** (47 mg, 111.5 µmol. 1.0 equiv.) was dissolved in DCM (750 µL), followed by the addition of TFA (250 µL) at 0 °C under stirring. The reaction mixture was removed from the ice bath and stirred for 1 h at RT. Excess TFA and DCM was removed in a Ar stream. Under cooling, the mixture was basified with 2 M aq. NaOH, extracted with 20% *i*PrOH in $CHCl_3$ (3 x 2 mL), briefly dried over $Na_2SO_4$, filtered and the solvent removed by rotary evaporation. The resultant freebase (25 mg, 111.5 µmol, 1.0 equiv.) was immediately dissolved in THF (3 mL), cooled to 0 °C, followed by addition of $Et_3N$ (31 µL, 223 µmol, 2.0 equiv.), then chloroacetic anhydride (19.1 mg, 111.5 µmol, 1.0 equiv.). The reaction was quenched by addition of TFA (1 equiv.), followed by rotary evaporation of all volatiles. The crude product was partially redissolved in 3 mL 5% aq. MeCN, 1 mL 50% aq. MeCN and 1 mL MeCN and then filtered over a 0.2 µm PVDF filter tip. The resultant clear solution was purified by injection into a preparative HPLC system (0 - 50% aq. MeCN over 30 min). The product was obtained as a white lyophilisate (7 mg, 23 µmol, 21%).

$^1$H-NMR (300 MHz, DMSO-$d_6$) δ ppm 14.1 (br s, 1H), 8.87 (d, *J* = 6.4 Hz, 3H), 8.39 (d, *J* = 6.4 Hz, 2H), 8.18 (s, 1H), 7.70 (d, *J* = 8.6 Hz, 2H), 7.46 (d, *J* = 8.6 Hz, 2H), 4.50 (d, *J* = 5.6 Hz, 2H), 4.15 (s, 2H). $^{13}$C-NMR (75 MHz, DMSO-$d_6$): δ ppm 166.5, 146.6, 145.7, 141.8, 139.2, 134.1, 127.6, 122.3, 121.0, 119.3, 112.0, 43.2, 43.2. LC-MS: m/z 301 corresponding to $[M+H]^+$, m/z 303 corresponding to $[M+H]^+$ for the heavier chloride isotope.

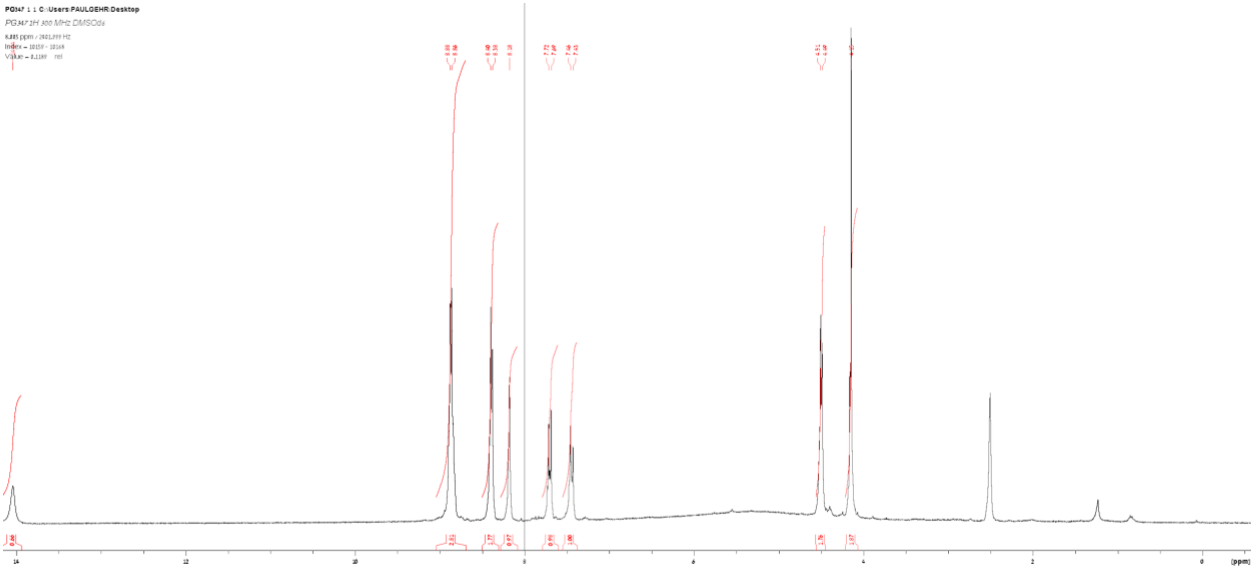

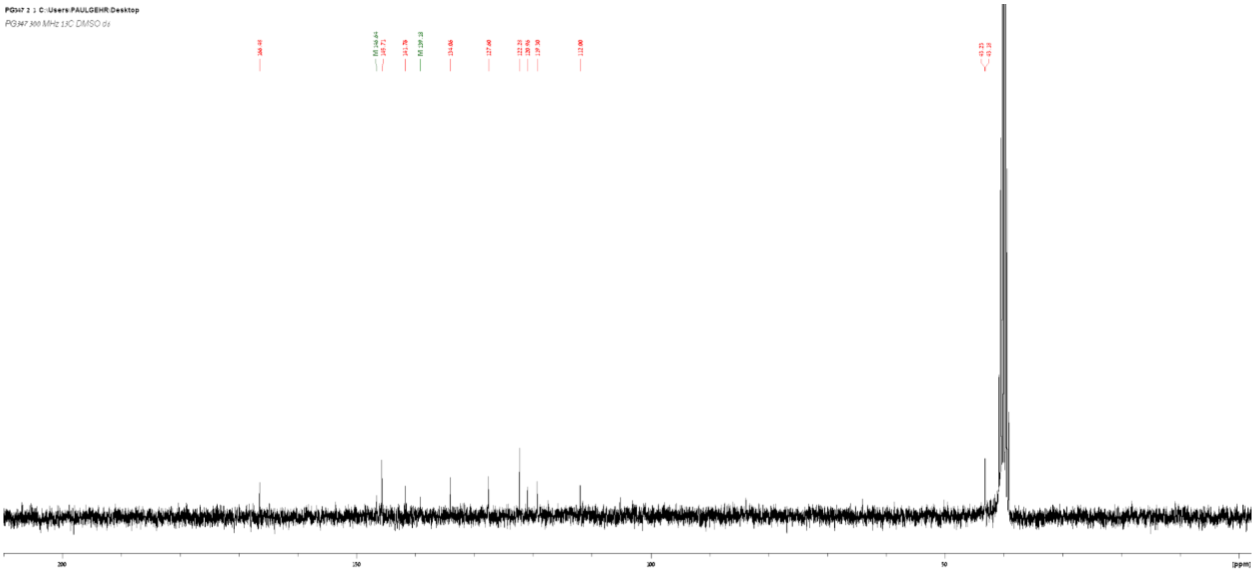

<u>*N*-(4-(tert-butyl)phenyl)-*N*-(2-(tert-butylamino)-2-oxo-1-(pyridin-3-yl)ethyl)acrylamide (**10**)</u>

A two dram vial with Teflon stir bar was charged with 1.5 mL MeOH (AR grade, non-dried), followed by the addition of pyridine-3-carbaldehyde (10.9 µL, 100 µmol), 4-tert-butylaniline (15.9 µL, 100 µmol), acrylic acid (7 µL, 100 µmol) and tBuNC (11.3 µL, 100 µmol) exactly in this order. The vial was capped, protected from light and the reaction mixture stirred for 24 h at RT. After checking complete conversion by LC-MS, the solvent was removed by rotary evaporation inside a fumehood. Reactionware contaminated with tBuNC was washed with 10% conc. HCl in MeOH (v/v). The crude product was dissolved in 50% aq. ACN and 100 µL DMF, filtered over a 0.2 µm PVDF or PTFE syringe tip frit and subsequently purified by preparative RP-HPLC. The product was obtained as a white, hygroscopic lyophilisate (24 mg, 61 µmol, 61%).

$^{1}$H-NMR (400 MHz, DMSO-$d_6$) δ ppm 8.65 - 8.60 (m, 2H), 8.01 (s, 1H), 7.89 (d, *J* = 8.1 Hz 1H), 7.60 (dd, *J* = 8.1 Hz, 5.3 Hz, 1H), 7.36 (d, *J* = 8.5 Hz, 2H), 7.22 (br s, 2H), 6.29 - 6.23 (m, 2H), 5.96 (dd, *J* = 16.8 Hz, 10.3 Hz, 1H), 5.66 (dd, *J* = 10.3 Hz, 2.0 Hz, 1H), 1.29 (s, 9H), 1.25 (s, 9H). $^{13}$C-NMR (101 MHz, DMSO-$d_6$): δ ppm 167.4, 165.2, 151.1, 147.3, 145.3, 142.1, 136.6, 134.6, 130.8, 129.3, 128.4, 126.0, 125.0, 62.1, 51.0, 34.7, 31.4, 28.7. LC-MS: m/z 394 corresponding to $[M+H]^+$. HRMS: m/z, $C_{24}H_{32}N_3O_2$ calculated 394.2495 $[M+H]^+$, found 301.2492 $[M+H]^+$, 0.8 ppm deviation.

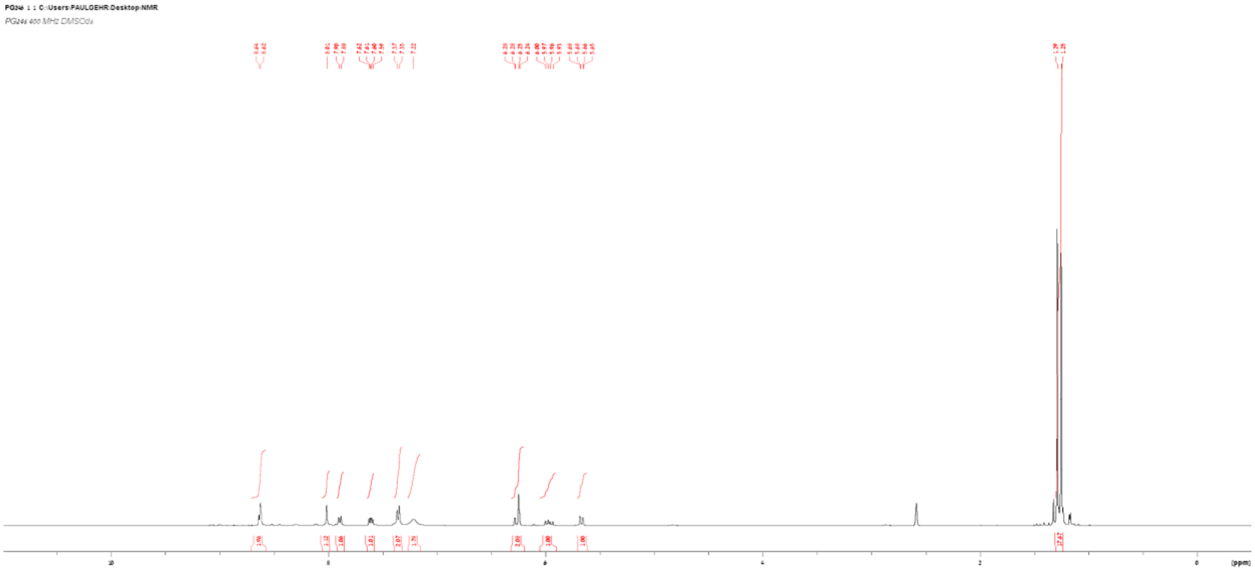

PG246 1 1 C:\Users\PAULGEHR\Desktop\NMR
PG246 400 MHz DMSOd6
10
8
6
4
2
0
[ppm]


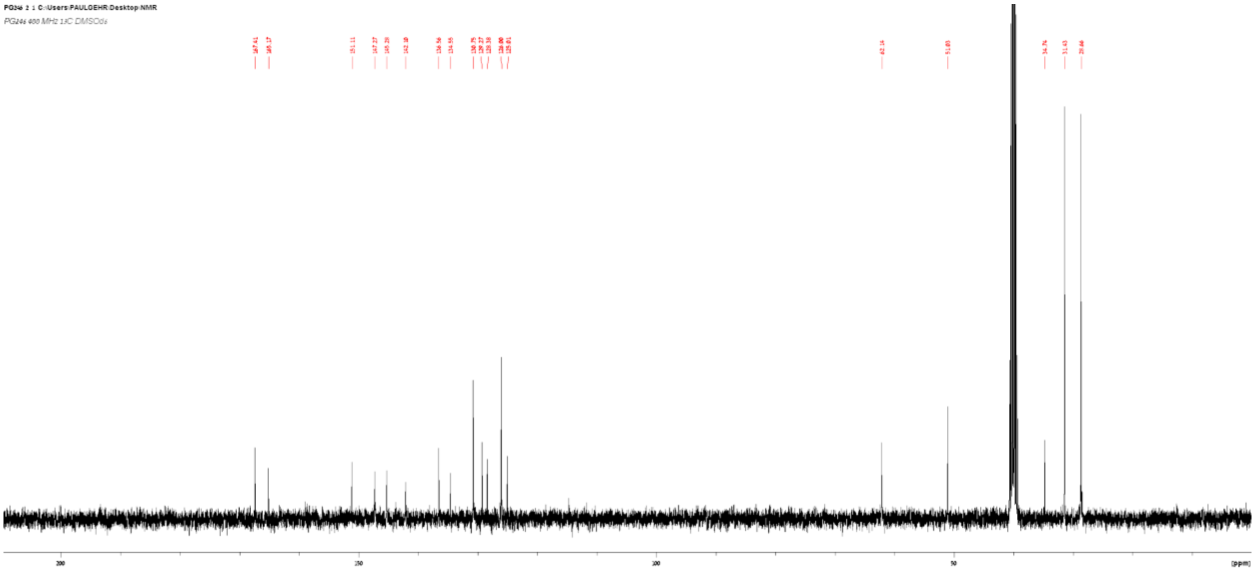

PG246 2 1 C:\Users\PAULGEHR\Desktop\NMR
PG246 400 MHz 13C DMSOd6
167.41
165.17
200
150
100
50
[ppm]
